# FGFR2 promotes resistance to ALK tyrosine kinase inhibitors and its inhibition acts synergistically with lorlatinib in the treatment of ALK-expressing neuroblastoma

**DOI:** 10.1101/2024.09.05.611416

**Authors:** Perla Pucci, Charlotte Barrett, Ricky Trigg, Jamie D. Matthews, Marcus Borenäs, Michaela Schlederer, Leila Jahangiri, Lucy Hare, Christopher Steel, Emily James, Nina Prokoph, Lukas Kenner, Ruth Palmer, Bengt Hallberg, G. A. Amos Burke, Suzanne D. Turner

## Abstract

Anaplastic Lymphoma Kinase inhibitors (ALK TKIs) are approved for the treatment of ALK-positive non-small cell lung cancer (NSCLC) and are in clinical trial for ALK-aberrant high-risk neuroblastoma (NB) patients, particularly loratinib. However, resistance to ALK inhibitors can occur in patients, via the activation of bypass-signalling pathways, and there is a need to identify these mechanisms as well as drugs that inhibit them to design therapeutic approaches that prevent resistance, and to treat ALK TKI relapsed/refractory disease. Using genome-wide CRISPR-Cas9 overexpression screens, we identified and validated *FGFR2* as a desensitizer to lorlatinib in aberrant ALK-expressing high-risk NB. FGFR2 and FGFR2-associated pathways are up-regulated in lorlatinib-resistant NB cells. Moreover, high-throughput screens using a library of 1,430 FDA approved drugs identified kinase inhibitors including those targeting FGFR2 as efficacious in reducing the survival of lorlatinib resistant NB cells. Hence, the FGFR pathway was investigated as a therapeutic target applying the pan-FGFR inhibitor erdafitinib or the multi-kinase inhibitor ponatinib, resulting in reduced survival of lorlatinib-resitant cells in comparison to their lorlatinib-sensitive counterparts. Moreover, both FGFR inhibitors act synergistically with lorlatinib *in vitro* and *in vivo*, using patient-derived xenografts (PDXs) and genetically engineered models (GEMM) of ALK-expressing NB. *FGFR2* mRNA expression also correlate with a poorer prognosis for NB patients, regardless of sub-type, suggesting that a broader range of patients may benefit from FGFR inhibitors. Overall, our data suggests that FGFR2 potentially plays roles in lorlatinib resistance in NB and that combined pharmacological inhibition of ALK and FGFR constitutes a therapeutic approach to treat high-risk NB.

## INTRODUCTION

Monotherapies can have limitations in maintaining durable responses without encountering the toxicities associated with high-doses, causing short– and long-term side effects^1–4^. Designing therapeutic strategies to target resistance mechanisms can help to treat relapsed disease or, improve progression free survival rates by preventing the emergence of resistance. Despite the initial benefit that anaplastic lymphoma kinase (ALK) tyrosine kinase inhibitors (TKIs) have shown, and the increasing number of successful trials using these agents for cancer treatment^3,5^, resistance can occur in patients and is associated with poorer survival in different cancers including neuroblastoma (NB). NB, originating during the aberrant embryonal development of neural crest cells, has the highest frequency and mortality of all extracranial solid cancers diagnosed in children and infants^6^. Most NB patients are diagnosed with high-risk NB (60%), which lacks effective treatments, resulting in a 5-year event-free survival (EFS) of less than 50%^7–12^. Among others, *MYCN* amplification (accounting for 20% overall and 50% of high-risk cases), *ALK* amplification and/or *ALK* kinase-domain point mutations (accounting for 8-10% overall and 15% of high-risk cases) are key molecular factors associated with aggressive phenotypes and poor treatment outcomes^8,9,13–16^. High-risk NB patients urgently need effective and kinder therapeutic approaches, of which ALK TKIs, and in particular lorlatinib has shown promising evidence^3,5^ but relapse is a distinct possibility as mentioned above^17–20^. Hence, additional drug targets for combination therapy of NB to prevent resistance and/or relapse are urgently needed^21–23^.

Several preclinical models of ALK TKI resistance show ALK TKI-dependent induction of bypass signalling revealing new vulnerabilities that can be targeted alongside ALK, to prevent or overcome resistance^24–26^. Therefore, combinatorial regimens targeting bypass signalling mechanisms are a promising treatment strategy for ALK-expressing NB and would also help achieve improved outcomes using lower individual drug doses in synergistic combinations, and therefore potentially also reduce toxicity^27^. Lorlatinib, the third-generation ALK TKI, is a promising drug for treating ALK-aberrant NB patients particularly as it has good central nervous system penetration^3,28,29^. Lorlatinib has also been shown to bind to ALK despite the presence of mutations associated with resistance to other ALK TKIs, and has increased binding affinity with the leu1198 residue within the ATP-binding pocket of ALK, thereby increasing its specificity and reducing the incidence of off-target cytotoxic effects^26,30,31^. However, despite this, lorlatinib is associated with side-effects, including weight gain, hyperlipidaemia and cognitive defects^23,32^. In addition, acquired resistance to lorlatinib has been demonstrated in model systems and patients^21,26,33^. One strategy to prevent and/or overcome the development of acquired resistance associated with ALK TKIs is a combinatorial therapeutic regimen allowing for lower doses of the single agents while still displaying superior efficacies over each drug acting alone^24–26,34^. Finding a drug that will act synergistically with ALK TKI treatment requires knowledge of the resistance mechanisms that develop in response to these drugs. As more potent ALK TKIs are utilised, the more pressing resistance mechanisms that develop will likely be ALK-independent^24,30^.

Pre-clinical studies support combination targeted therapy as a viable treatment strategy for ALK-driven NB^24–26,34–37^. Combination therapies such as lorlatinib and MEK inhibitors have been investigated for NB treatment showing efficacy for lorlatinib-resistant NB cells with NF1 loss, but not in those that acquired the NRASQ61K mutant, suggesting that this combination could be clinically relevant for a subset of ALK TKI resistant NB patients^26^. Additionally, the MEK inhibitor Trametinib has shown limited efficacy in ALK-expressing NB due to activation of pro-survival feedback signalling through Akt/mTORc^38^. Lorlatinib has also been assessed for efficacy when used together with the MDM2/MDMX inhibitor idasanutlin in NB PDXs where this combination resulted in significant inhibition of tumour growth in one of three models which carried an ALK amplification, but no response in the PDX carrying an ALK mutation^34^. These findings highlight the complexity of signalling pathways and the challenge to find a collateral sensitivity common to the majority of ALK expressing and/or ALK TKI resistant NB patients.

With this aim, we performed genome-wide CRISPR-dCas9 overexpression screens (CRISPRa) in two distinct cell lines of aberrant ALK-expressing NB, to identify genes involved in the response to lorlatinib. These screens identified FGFR2, which when upregulated *in vitro*, desensitized cells to lorlatinib. RNA sequencing (RNA-Seq) of lorlatinib-resistant NB cells (LR), including our newly developed models, was conducted to find potential bypass resistance tracks to lorlatinib, identifying FGFR2 expression and signalling as the most represented pathway. Furthermore, high-throughput drug screens of the lorlatinib-resistant NB cells and their lorlatinib-naïve counterparts showed that tyrosine kinase receptors were a common target among drugs that were significantly more efficacious in the LR than in the parental cell lines, including inhibitors targeting the FGFR pathway. FGFR2 was therefore further investigated as a therapeutic target using the FDA-approved FGFR inhibitor erdafitinib and the multi-kinase inhibitor ponatinib. We show that both drugs act synergistically with lorlatinib, decreasing cell viability of both lorlatinib-naïve parental and LR NB cell lines, and *in vivo* PDX models. Furthermore, we show that FGFR2 mRNA expression correlates with poor overall survival (OS) and EFS of NB patients, suggesting that FGFR inhibitors may have broader applicability to the treatment of NB. Overall, our data suggest that FGFR2 plays a role in the cellular response to ALK TKIs and that combined pharmacological inhibition of ALK and FGFR constitutes a therapeutic approach to treat high risk ALK-aberrant NB.

## RESULTS

### Genome-wide CRISPRa screens identify FGFR2 as a potential therapeutic vulnerability in ALK-aberrant NB treated with the ALK TKI lorlatinib

Genome-wide CRISPRa screens were conducted of the dCas9-expressing NB cell lines SH-SY5Y (ALK^F1174L^) and CHLA20 (ALK^R1275Q^) following exposure to the ALK TKI lorlatinib, at the effective concentration killing 50% of the cells (EC_50_) for 14 days. Prior to drug exposure, the dCas9 expressing cells were transduced with an sgRNA library containing 3 sgRNA/gene, targeting 23,430 genes^39,40^. Genomic DNA was extracted from the cells at day 0 and at day 14 of drug exposure and next-generation sequencing (NGS) was performed to identify genes, which following their overexpression, allow aberrant ALK-expressing NB cells to survive in the presence of lorlatinib, therefore representing potential bypass mechanisms (Supplementary Fig. 1 left panel).

Exposure of CHLA20 or SHSY5Y cells to lorlatinib (500 nM) led to significant enrichment of gRNAs targeting 254 and 391 genes respectively (beta score>1; *p*<0.01, permutation-based non-parametric analysis (PBNPA)^41^) (Fig. 1A, B, Supplementary Fig. 2A, Extended Data File 1). *FGFR2, MET* and *BACH2* were significantly enriched in both cell lines although gene set enrichment analysis (GSEA) of candidate resistance genes detected by the screens in both cell lines, in comparison with ‘hallmark of cancer and canonical pathway gene sets’ in the Molecular Signatures Database^42^, showed that FGFR2 was the only gene detected in both cell lines and represented in the ‘top 10 most representative of cancer pathways’ (Fig. 1C, Supplementary Fig. 2C). Furthermore, pathway analysis using Reactome revealed that most of the pathways commonly upregulated in both lines are FGFR2-associated pathways (Fig. 1D, Supplementary Fig. 2B).

**Figure 1.**
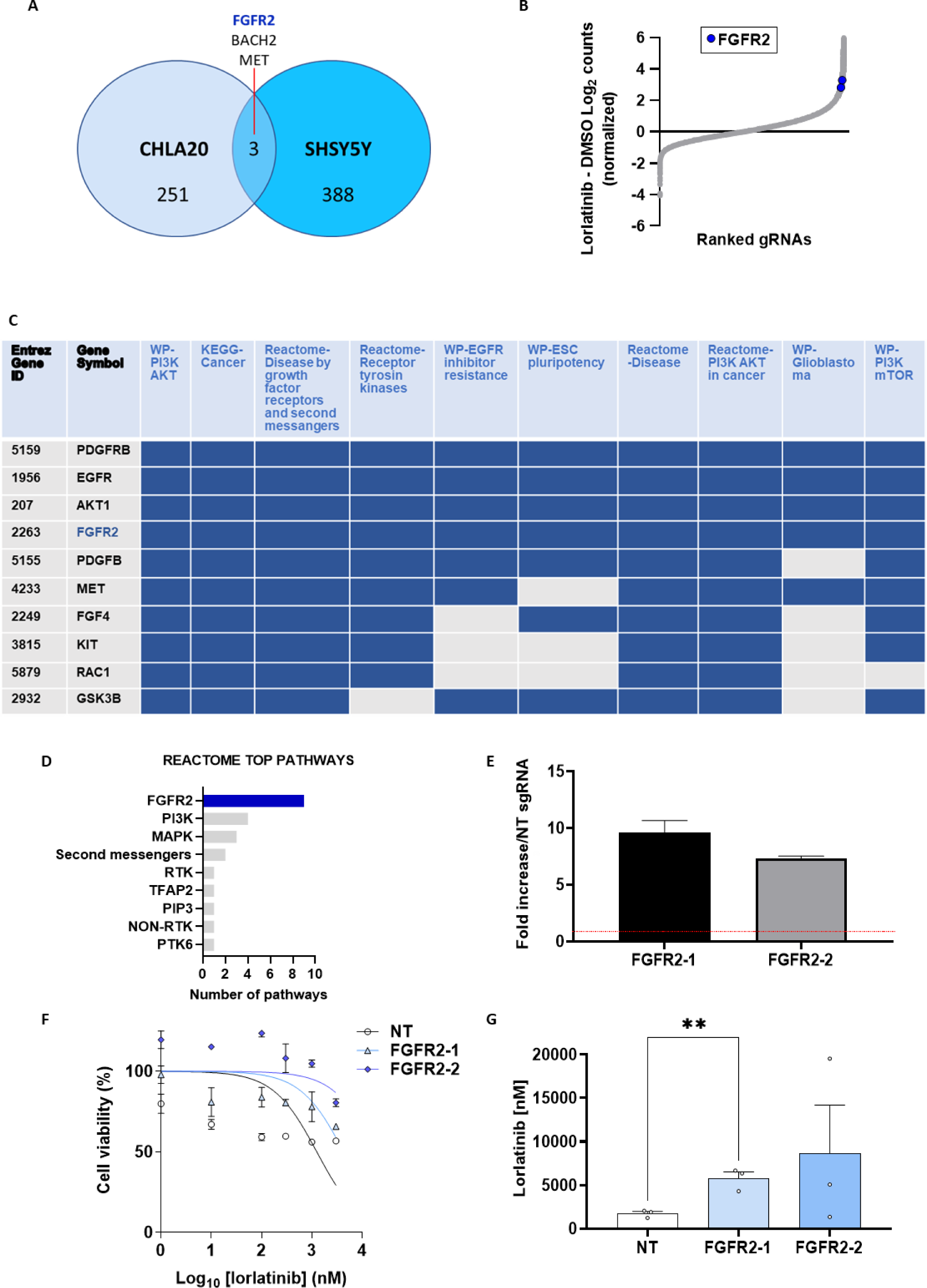
| CRISPRa screens identify FGFR2 expression as a desensitizer to lorlatinib treatment. (**A**) FGFR2, BACH2 and MET were the only commonly enriched hits identified by CRISPRa screens conducted in both cell lines (CHLA20 and SH-SY5Y) with a beta score>1; *p*<0.01. (**B**) Significantly enriched genes (beta score>1, p<0.01, permutation-based non-parametric analysis (PBNPA)^41^) from CRISPRa screens conducted in CHLA20 cells. The gRNAs are ranked by fold-change vs difference in log-normalized read counts between DMSO and lorlatinib-treated CHLA20 cells. Two gRNAs per gene are shown and the gRNA to FGFR2 is highlighted in blue. (**C, D**) FGFR2 and its regulated pathways are enriched in CHLA20 NB cells following 14 days of lorlatinib treatment compared to DMSO treated cells: GSEA with GO gene sets in MSigDB in C and Reactome in D. (**E**) FGFR2 expression levels in CHLA20-dCas9 cells transduced with one of two independent FGFR2 gRNAs (FGFR2-1 and FGFR2-2) compared to non-targeting (NT) gRNA transduced cells. (**F**) Viability (cell-titer blue (CTB)) of CHLA20-Cas9 cells transduced with NT gRNA or two different FGFR2 gRNAs following treatment with the indicated doses of lorlatinib for 72 h. (G) EC_50_ values calculated from (F) are shown as means ±SEM from three biological replicates. Significance was determined using a two-tailed Student’s t-test, **p=0.0066.

### Overexpression of FGFR2 in neuroblastoma cell lines decreases their sensitivity to the ALK TKI lorlatinib

Candidate resistance genes identified by the CRISPRa screens conducted in SH-SY5Y and CHLA-20 cell lines were each functionally validated by transducing Cas9-expressing cells with each of two gene-specific gRNAs individually and determining cell viability in response to lorlatinib exposure (Fig. 1E-G, Supplementary Fig. 2 D-E). Given the consistent identification of FGFR2 and FGFR2-mediated pathways, we focussed our screen validation on FGFR2. FGFR2 overexpression was achieved using both FGFR2-specific gRNAs for both cell lines as assessed by RT-qPCR (Fig. 1E, Supplementary Fig. 2D). Furthermore, FGFR2 overexpression led to an increase in cell viability under lorlatinib treatment (Fig 1F, Supplementary Fig 2E) with a subsequent increase in the lorlatinib EC_50_ (Fig. 1G).

### FGFR2 is upregulated in cell line models of acquired resistance to lorlatinib

In order to confirm whether FGFR2 overexpression represents a *bone fide* bypass signalling mechanism in NB, lorlatinib resistant NB cell lines (KELLY-LR, CHLA20-LR and CLBGA-LR^21^), generated following exposure to serially increasing concentrations of lorlatinib over 6 months (Supplementary Fig. 1 middle panel, Supplementary Fig. 3A) were analysed by bulk RNA-seq for differential gene expression (DE). Analysing 4 biological replicates of each cell line, no change in the status of exonic *ALK* mutations considered deleterious and damaging for the protein, were detected in LR compared to parental lines, suggesting further mutations in ALK were not responsible for the observed lorlatinib resistance (Extended Data File 2). As expected, principal component analysis (PCA) and a correlation matrix heatmap of the RNA-seq data showed that cell lines of the same origin clustered together (Supplementary Fig. 3B-C). Genes significantly upregulated showed a large variation in log_2_ fold change (Fig. 2A, Supplementary Fig. 3D). The top genes upregulated in LR compared to parental cell lines (with the highest base mean >100 for individual samples and >500 as the average for all cell lines, a fold change of log_2_>1.5 and *p*<0.05) include *FGFR2, FAT1, SPOCK1, ERBB4, EGFR, PDE3A, TSHZ2, CACNA1D, COL5A1, COL18A1, ADCYAP1, TNFRSF19, APOE* and *PIEZO1* (Fig. 2A).

**Figure 2.**
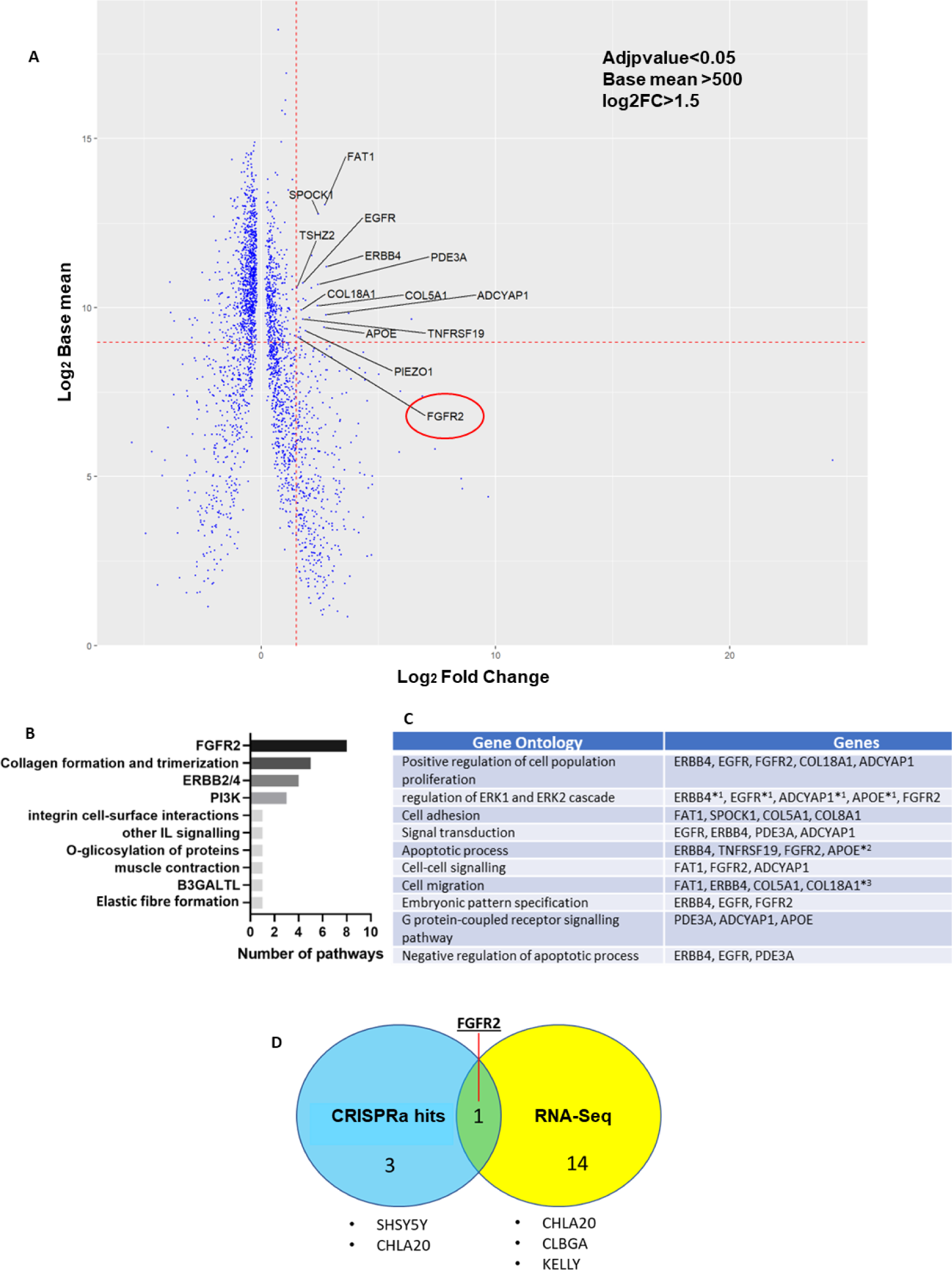
| FGFR2 is up-regulated in NB cell lines with acquired resistance to lorlatinib. (**A**) A 2D Plot of log2 fold change (FC) against the –log10 of P adjusted values (Adjp) of RNA for all cell lines (CHLA20, CLBGA and KELLY versus their LR counterparts). Log2 FC>0 = genes with higher expression in LR cell lines. FGFR2 (highlighted by a red circle) is one of the 14 top upregulated genes. (**B**) The top 25 pathways from Reactome analysing all significantly upregulated genes detected in the LR cells compared to their respective parental counterparts (CHLA20, KELLY and CLBGA), and grouped according to the common main pathway regulator. (**C**) The 10 most frequently occurring Gene Ontology (GO) terms of biological processes associated with the 14 significant genes (including FGFR2) upregulated in LR cell lines. (**D**) Diagram showing hits from the CRISPRa screens significantly enriched in both cell lines used (SHSY5Y and CHLA20) merged with the identified 14 genes upregulated in LR cells compared to their respective parental cells (CHLA20, KELLY and CLBGA), determined by RNA-Seq.

In keeping with these data, pathway overrepresentation analysis using the REACTOME database^43^ showed that among the top 25 overrepresented pathways associated with the 599 significantly (*p*<0.05) overexpressed genes, those involved in FGFR2 activation and signalling were the most prevalent for all 3 cell line pairs (Fig. 2B). Furthermore, GO analysis of the 14 top genes that were identified to be putative lorlatinib resistance drivers showed an overarching theme of ‘signal transduction’ and three additional biological processes: ‘proliferation’, ‘apoptosis’ and ‘migration’, suggestive of the biological processes that can be compensated for under lorlatinib treatment (Fig. 2C). Of note, FGFR2 was prevalent amongst these processes being associated with ‘proliferation’, ‘MAPK signalling’, ‘apoptosis’, ‘cell-cell signalling’ and ‘embryonic pattern specification’ processes (Fig. 2C). Hence, expression of FGFR2 was assessed by western blot in the LR cell lines as well as their parental counterparts revealing specific upregulation of FGFR2 in the LR cell lines (Supplementary Fig. 2E). These data are in keeping with the CRISPRa screen data whereby FGFR2 was amongst the 3 genes upregulated in both lorlatinib treatment-surviving cell lines and the 14 genes in the LR cell lines (Fig. 2D). Together, these data suggest that FGFR2 may play a key role in regulating lorlatinib response bypass pathways.

### High-throughput screens of FDA-approved drugs reveal novel vulnerabilities of lorlatinib-resistant NB cells

A quality controlled, high-throughput screen performed in four NB cell lines (KELLY, CHLA-20 and their LR counterparts) exposed to a library of 1,430 FDA approved drugs was conducted (Supplementary Fig. 1 right panel, Supplementary Fig. 4A-C). To select drugs specifically targeting resistance bypass mechanisms, we first considered drugs that did not decrease cell viability below 50% for parental cell lines but did for the LR counterparts (Fig. 3A, Supplementary Fig. 4D-E, Supplementary Fig. 5A). We also noted that several drugs that reduced cell growth by 50% or more for the LR cell lines, also caused a greater than 50% decrease in viability in the corresponding parental cell line, and that some drugs had a greater anti-tumorigenic effect on the parental cell line compared to the LR cells (Fig. 3A, Supplementary Fig. 4D-E, Supplementary Fig. 5A).

**Figure 3.**
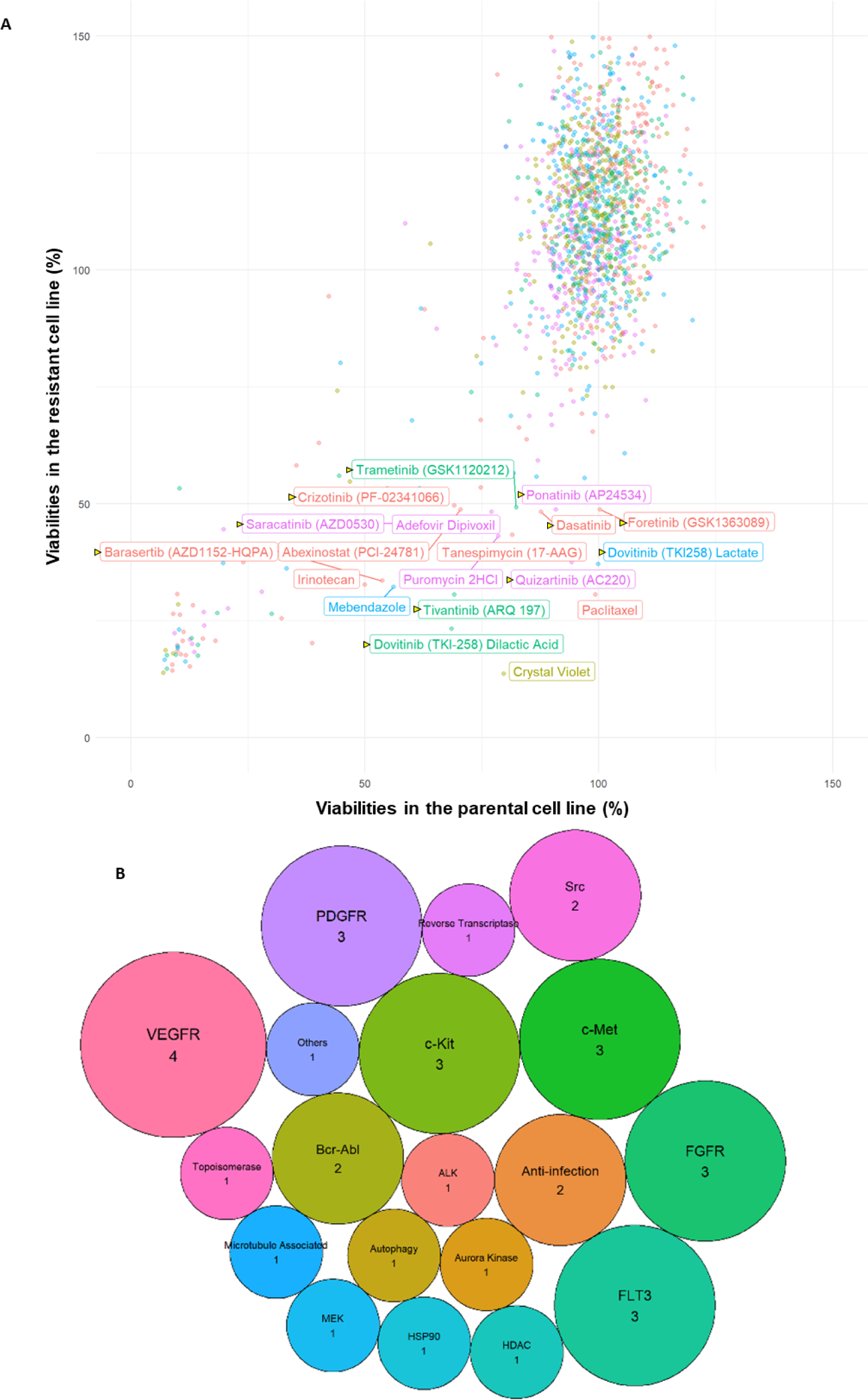
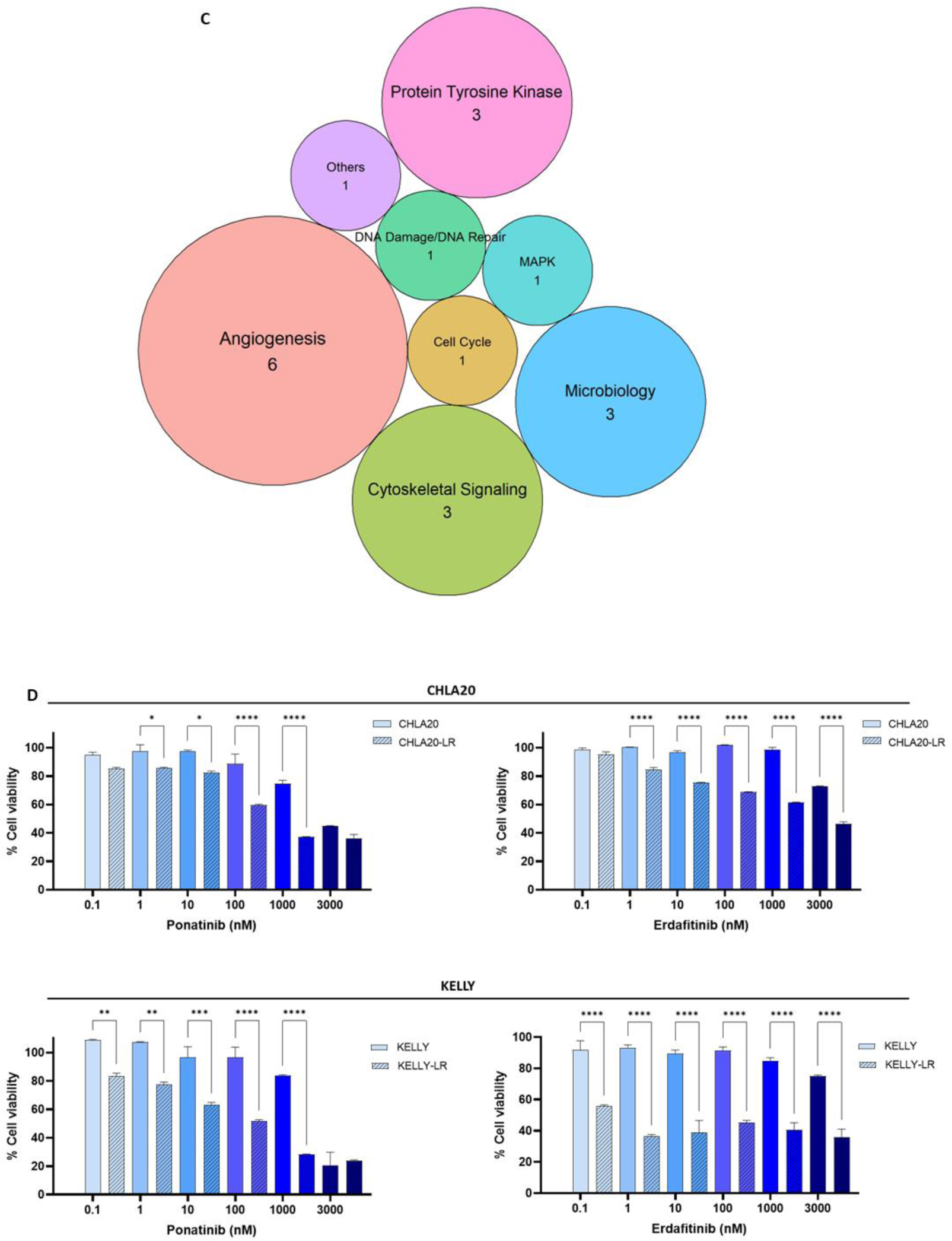
| High-throughput FDA-approved drug screens in CHLA20 and CHLA20-LR cells. (**A**) High-throughput FDA-approved drug (∼1500 drugs) screens performed with CHLA20 and CHLA20-LR cell lines showing cell viabilities following drug treatment (all drugs at 1000nM) for 72h. Those highlighted are the drugs that reduced viability of the LR cells to below 50% without similarly affecting the parental cells. The colours represent individual 384-well plates according to where the drugs are located in the library. Yellow triangles next to drug labels highlight the drugs inhibiting kinases. (**B**) Bubble plots showing the common targets for drugs more effective in killing CHLA20-LR cells than the parental cell line. (**C**) Bubble plots showing the common families of drugs (including protein tyrosine kinase) more effective in reducing the viability of CHLA20-LR cells than the parental cell line. (**D**) Cell viability (determined by CTB assay) in parental or LR cells (CHLA20 and KELLY) upon treatment with targeted inhibitors (ponatinib or erdafitinib) across a range of concentrations (0.1-3000nM), for 72h. Significance was determined using a one-way ANOVA with Sidak’s multiple comparison test of the means ± SEM from two biological replicates. *p<0.05, **p<0.01, ***p<0.001, ****p<0.0001. Above left panel: *p=0.011, 0.048, ****p= 0.000035, 0.0000026; above right panel: ****p= 0.000000023540776, 0.000000000154560, 0.000000000038229, 0.000000002162379, 0.000000735962777; below left panel: **p=0.0049, 0.00136, ***p= 0.00046, ****p= 0.000029, 0.0000029; below right panel: ****p= 0.0001, 0.0000029, 0.0000079, 0.000012, 0.000044, 0.00000086.

Among the 1,430 drugs tested, 19 were identified as efficacious in reducing cell viability below 50% for LR cells whilst not similarly affecting the parental CHLA-20 cell line, and similarly 13 drugs for the KELLY cell line (Fig. 3A, Supplementary Fig. 5A, Extended Data File 3). The known targets for the majority of the identified drugs are kinases including FGFR (highlighted by yellow arrows; Fig. 3A,B, Supplementary Fig. 5A,B). The shared drug targets correspond to common pathways: cytoskeletal signalling, angiogenesis, protein tyrosine kinase and microbiology (Fig. 3C, Supplementary Fig. 5C), suggesting a potential overlap in resistance bypass mechanisms utilised by each cell line, despite having heterogeneous characteristics, such as different ALK mutations, and differences in *TP53* and MYCN status.

Notably, drugs that target FGFR proteins were more effective at reducing the viability of both CHLA20-LR and KELLY-LR cells, but some were not observed in Fig 3A and Supplementary Fig. 5A due to the 50% median cut-off used (Table 1). Due to the specific identification of FGFR2 overall in our datasets, a selective FGFR inhibitor which was not included in the drug library was assessed for its efficacy in reducing cell survival. In addition, considering all data combined, the multi-kinase inhibitor ponatinib, already approved for use in children and targeting FGFR as well as a plethora of other kinases (BCR-ABL, FLT3, c-KIT, VEGFR, PDGFR and c-SRC^44^) was also assessed. Both drugs led to a reduction in viability of both LR cell lines and their lorlatinib-sensitive counterparts in a drug concentration-dependent manner with the LR cells, being significantly more sensitive as predicted by the drug screens (Fig. 3D).

**Table 1:** NB cell viability in response to 72hrs exposure to 1000nM of select FGFR or multi-kinase inhibitors that also target FGFR. LR = Lorlatinib Resistant.

| Drug | Targets | Median cell viability (%) |  |  |  |
| --- | --- | --- | --- | --- | --- |
|  |  | CHLA20 | CHLA20-LR | KELLY | KELLY-LR |
| **Erdafitinib** | FGFR 1-4 | 90.75 | 65.06 | 75.30 | 39.85 |
| **Infigratinib** | FGFR 1-3 (4) | 94.58 | 71.86 | 88.90 | 109.43 |
| **Pemigatinib** | FGFR 1-3 (4) | 93.08 | 79.78 | 97.18 | 96.1 |
| **Futibatinib** | FGFR 1-4 | 92.09 | 71.05 | 88.02 | 94.77 |
| **Ponatinib** | Abl, PDGFRα, VEGFR2, FGFR1-4, Src | 90.88 | 48.78 | 88.81 | 74.28 |
| **Ienvatinib,** | VEGFR1/Flt-1, VEGFR2/3, PDGFRα/$\beta$,<br>FGFR1-4, PDGFR, c-Kit, c-RET | 99.01 | 109.38 | 79.5 | 107.19 |
| **Regorafenib** | VEGFR1, VEGFR2, VEGFR3, PDGFRβ, Kit<br>(c-Kit), RET (c-RET) and Raf-1 | 105.80 | 123.16 | 87.36 | 115.10 |
| **Pazopanib** | VEGFR1, VEGFR2, VEGFR3, PDGFRβ, c-Kit,<br>FGFR1, and c-Fms/CSF1R | 104.44 | 84.75 | 91.90 | 97.28 |
| **Derazantinib** | FGFR 1-4, RET, DDR2, PDGFRβ, VEGFR<br>and KIT | 89.67 | 69.09 | 93.23 | 105.64 |
| **Dovitinib**<br>**dilactic** | FLT3/c-Kit, FGFR1/3, VEGFR1-4, InsR,<br>EGFR, c-Met, EphA2, Tie2, IGFR1, HER2 | 68.53 | 23.28 | 94.39 | 40.08 |
| **Dovitinib**<br>**lactate** | FLT3/c-Kit, FGFR1/3, VEGFR1-4, InsR,<br>EGFR, c-Met, EphA2, Tie2, IGFR1, HER2 | 99.92 | 37.10 | 74.31 | 39.57 |
| **Nintedanib** | VEGFR1/2/3, FGFR1/2/3 and PDGFRα/$\beta$ | 101.65 | 68.72 | 68.20 | 59.01 |
| **Dovitinib** | FLT3/c-Kit, FGFR1/3, VEGFR1-4, InsR,<br>EGFR, c-Met, EphA2, Tie2, IGFR1, HER2 | 100.79 | 87.0 | 93.49 | 118.79 |

### A combination of FGFR and ALK TKIs act synergistically in aberrant ALK-expressing NB cell lines

Given the activity of the inhibitors as single agents on both LR and parental cell lines, synergy between FGFR (ponatinib and erdafitinib) and ALK inhibitors (lorlatinib) was assessed. Applying a range of concentrations of ponatinib or erdafitinib with lorlatinib showed high synergism across multiple cell lines of both parental and LR origins, although the LR cells, in general, were more sensitive, confirming prior observations (Fig. 3D, Fig 4). The synergy experiments showed that ponatinib, in the absence of lorlatinib, was more efficacious in reducing viability of the KELLY-LR than the parental KELLY cell line at most concentrations whereby a reduction in cell viability is seen with as little as 0.1nM ponatinib (compared to 1000nM required for the parental cells) (Fig. 4A, Supplementary Fig. 6A) and for CHLA20-LR at concentrations greater than 100nM (again, compared to 1000nM for the parental cell line) (Fig. 4A, Supplementary Fig. 6A). The specific FGFR inhibitor erdafitinib (in the absence of lorlatinib) also shows greater efficacy for the LR cell lines although erdafitinib is significantly more potent for KELLY-LR compared to CHLA20-LR (Fig. 4A, Supplementary Fig. 6A). Indeed, erdafitinib reduced cell viability when given as a single agent to KELLY-LR cells at very low doses; less than 50% viability was seen following exposure to just 0.1nM.

**Figure 4.**
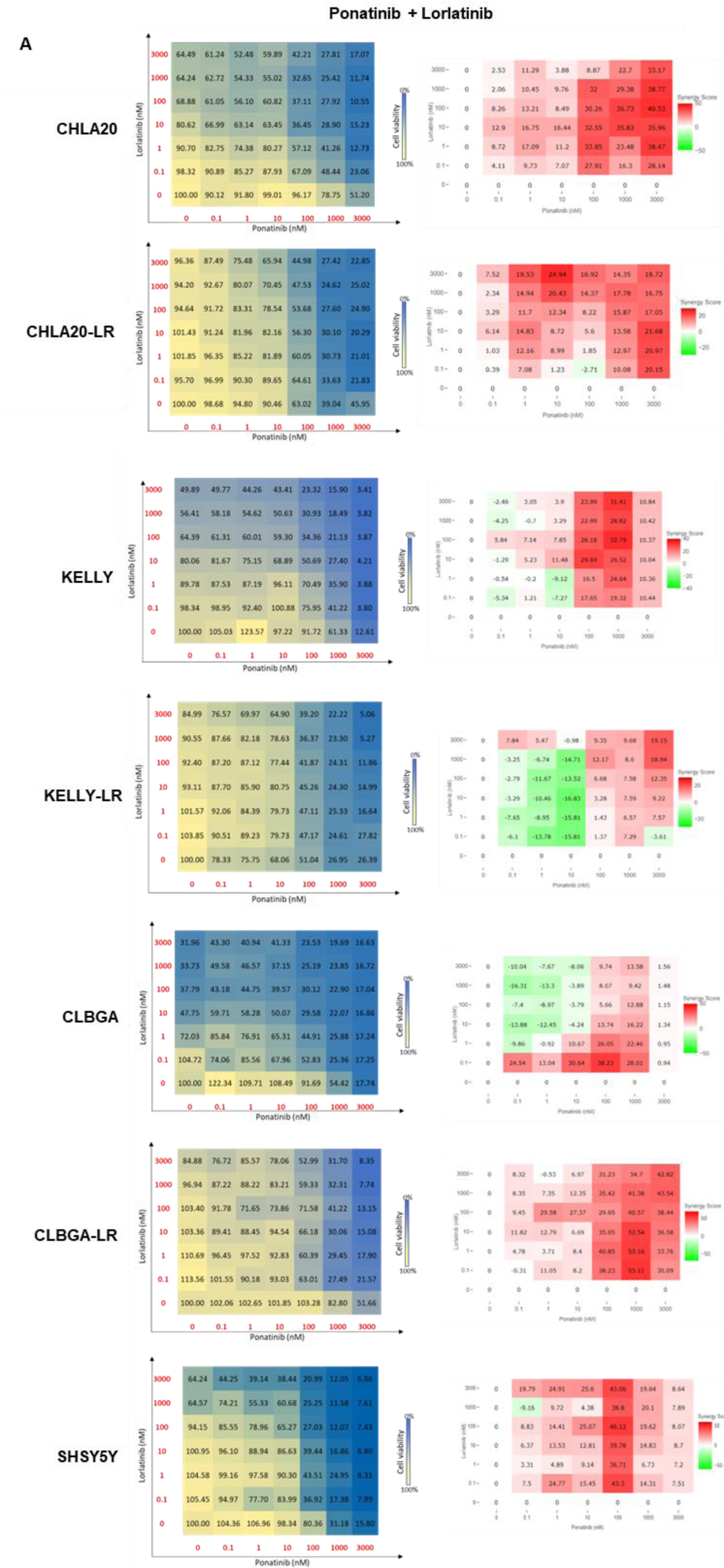
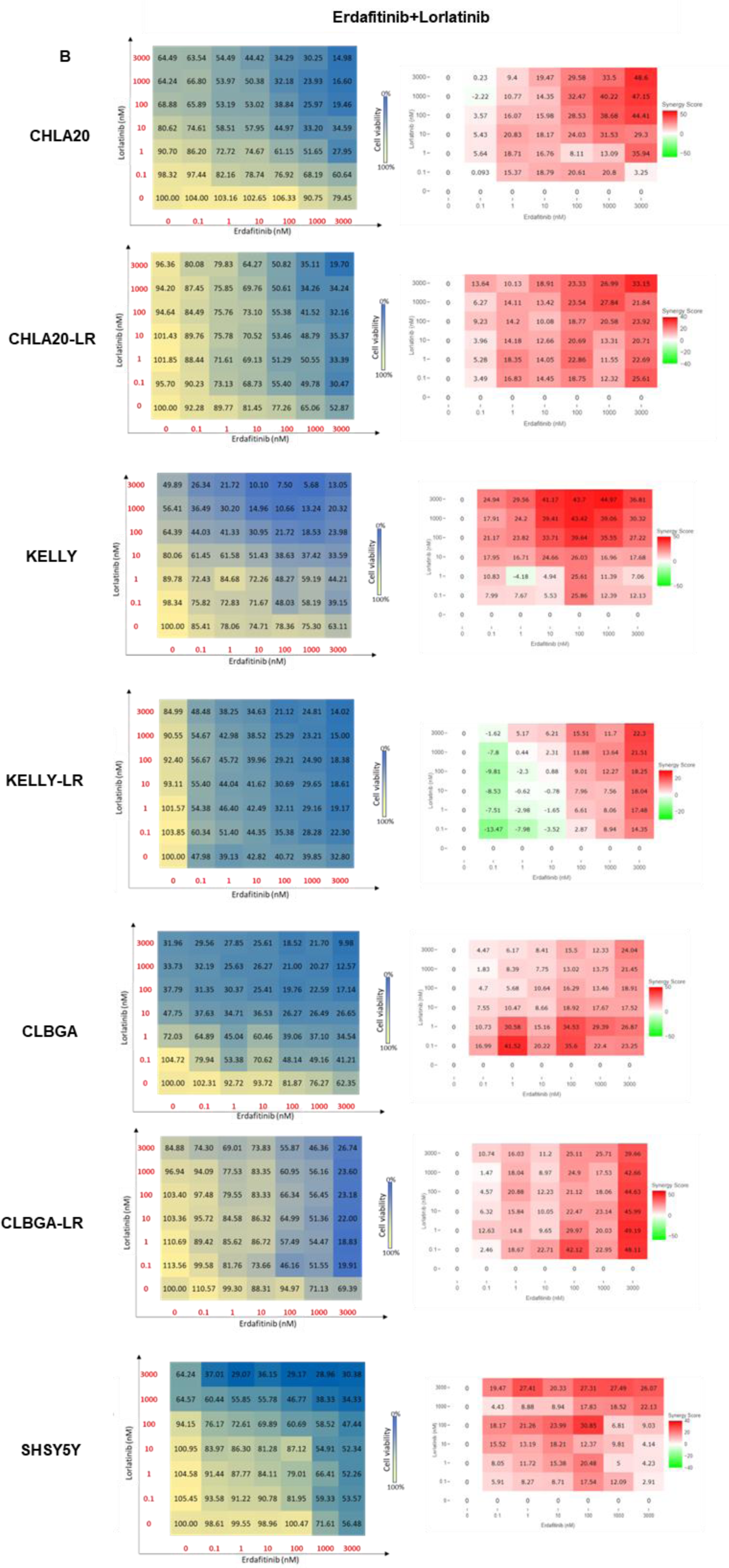
| Pharmacological inhibition of FGFR is synergistic with lorlatinib in NB cell lines and their LR-derived counterparts. (**A**) Dose-response matrix of ponatinib (0.1-3000nM) and lorlatinib (0.1-3000nM) alone or in combination, following 72 hrs incubation with the indicated cell lines. Panels on the left show cell viability; blue to yellow colour gradients: % cell viability normalised to DMSO treated cells (from yellow: 100%, to blue: 0%). Panels on the right represent the Loewe synergy scores; green to red colour gradients: synergy scores either with antagonistic or no effects (green, scores <-10) or synergistic (red, scores >10); scores between –10 and 10, represent additive effects. (**B**) Dose-response matrix of erdafitinib (0.1-3000nM) and lorlatinib (0.1-3000nM) alone or in combination, following 72 hrs incubation with the indicated cell lines. Panels on the left show cell viability; blue to yellow colour gradients: % cell viability normalised to DMSO treated cells (from yellow: 100%, to blue: 0%). Panels on the right represent the Loewe synergy scores; green to red colour gradients: synergy scores either with antagonistic or no effects (green, scores <-10) or synergistic (red, scores >10); scores between –10 and 10, represent additive effects. Results represent two biological replicates for each of the seven cell lines assessed.

In general, erdafitinib showed higher synergy scores than ponatinib when given in combination with lorlatinib across cell lines regardless of MYCN status (MYCN amplified KELLY or non-amplified CHLA-20 cell lines) (Fig. 4A, B, Supplementary Fig. 6A, B). These findings were confirmed for CLBGA, CLBGA-LR and SHSY5Y cell lines. Notably, cells highly sensitive to at least one of the two agents alone, such as CLBGA or KELLY-LR, showed lower synergy scores, suggesting that cell lines showing reduced efficacy to the drugs when administered as single agents, might benefit more from this combination approach (Fig. 4A, B, Supplementary Fig. 6A, B). While the effects of the FGFR inhibitors given as sole agents, in general, were higher for LR cells, suggesting the importance of FGFRs as a bypass mechanism for lorlatinib resistance, combination treatments were also highly effective in these cells, suggesting that FGFR inhibitors could re-sensitise cells to lorlatinib. Hence, this evidence supports the use of these combinations as not only upfront therapeutic approaches for treatment-naïve disease to prevent resistance from occurring in the first place, but also for relapsed/refractory/resistant cases.

### ALK TKIs and FGFR inhibitors show synergistic activity in the treatment of NB patient derived xenografts and GEMM *in vivo* models

Due to the noted effect of ALK TKI combination treatments with FGFR inhibitors in NB cells, we further investigated this regulatory axis in the ALK mutant, high-risk NB PDX, COG-N-415 (ALK_F1174L_; MYCN amplified) and COG-N-496x (ALK WT; MYCN amplified). NSG mice were injected subcutaneously with early passage NB PDX cells suspended in Matrigel, and when the tumours reached approximately 100mm^3^ the mice were treated daily with either vehicle (20% hydroxypropyl-beta-cyclodextrin), low dose single-agent lorlatinib (10 mg/kg), single-agent erdafitinib (30 mg/kg) or both agents in combination at the same doses (n=7) (Fig. 5A-D). Erdafitinib was selected over ponatinib for *in vivo* testing since it specifically targets FGFRs, while also showing promising effects when used in combination with lorlatinib *in vitro* (Fig. 4). Single-agent treatment with sub-optimal concentrations of lorlatinib or erdafitinib partially delayed tumour growth compared to vehicle alone (Fig 5A,B; corresponding to a median EFS of 18 days, and 17 days, for single agent lorlatinib or erdafitinib respectively, versus 11 days for untreated COG-N-415x tumour-bearing mice (Fig. 5C) and to a median EFS of 24 days for lorlatinib versus 16 days for untreated COG-N-496x (Fig. 5D); although no difference was observed using erdafitinib alone). However, combination drug treatment with lorlatinib and erdafitinib led to remissions with significant reductions in tumour volume (p<0.0001) and a corresponding significant increase in EFS (p<0.01 for COG-N-415x and p<0.05 for COG-N-496x) for all of the animals (Fig. 5A-D). All compounds were well-tolerated, with no significant decrease in body weight nor lethal toxicity observed (Fig. 5E, F).

**Figure 5.**
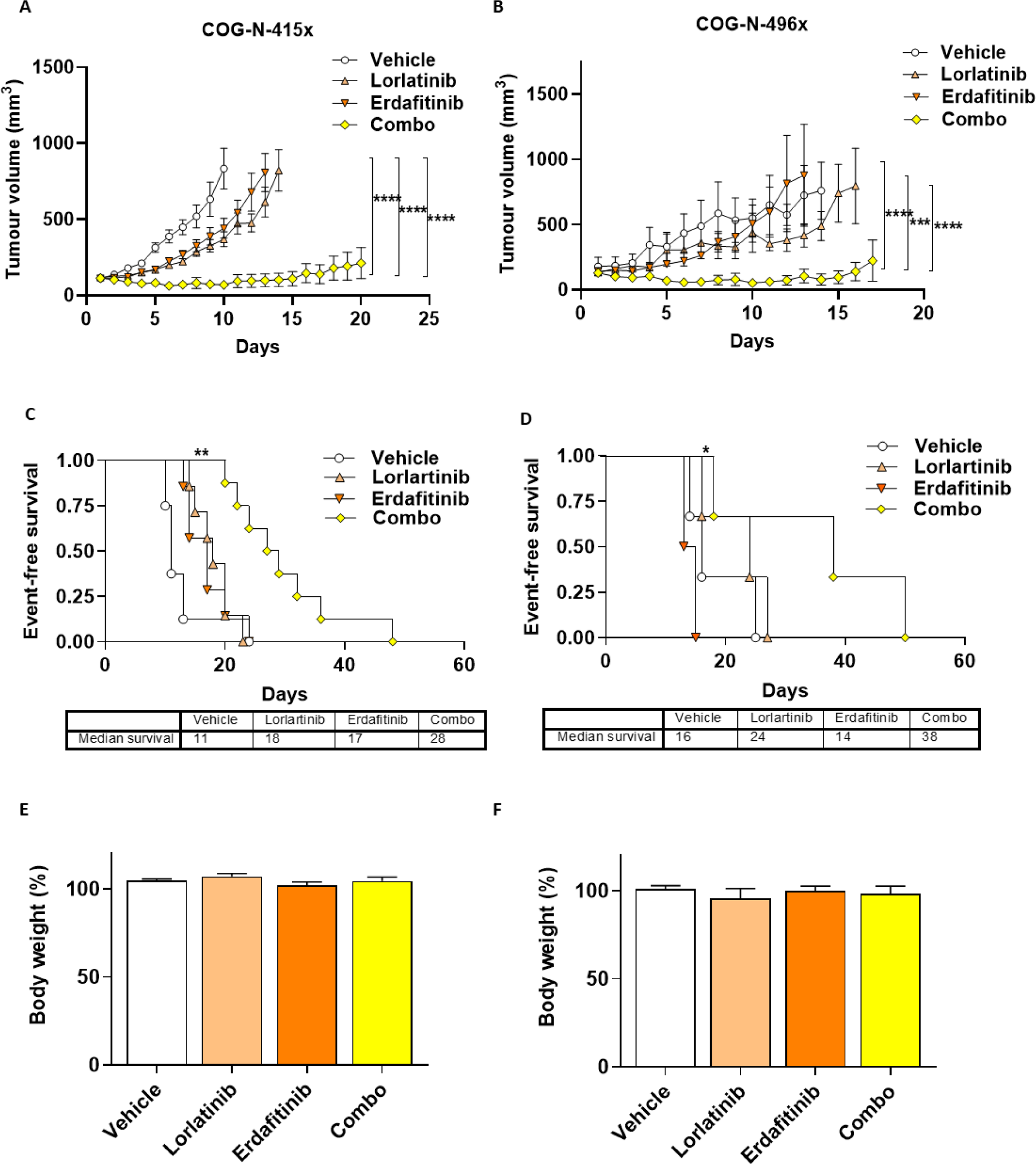
| A combination lorlatinib with the FGFR inhibitor erdafitinib significantly reduces tumour growth *in vivo* in PDX models of NB. (**A, B**) Tumour volume over time of NSG mice injected sub-cutaneously with COG-N-415x (A) or COG-N-496x (B) primary NB cells which reached 100mm^3 before daily administration of either vehicle (20% hydroxypropyl beta cyclodextrin), lorlatinib (10 mg/kg), erdafitinib (30 mg/kg), or lorlatinib and erdafitinib (combo, same doses). The study endpoint is tumours reaching 15 mm diameter or following 21 days of treatment, whichever came first. Data shown represent means ± SEM from 7 mice (n = 7) at each time point. Significance was determined using a one-way ANOVA with Tukey’s post-test at each experimental endpoint. ****p = 0.0000059, 0.000000000000077 and 0.00000000077 respectively (A) and ***p= 0.001, ****p =0.000010 or 0.000070 respectively (B). (**C, D**) Kaplan–Meier EFS analysis for mice carrying either COG-N-415x (C) or COG-N-496x (D) cells. Data points (n = 7) represent means ± SEM, shown until the experimental endpoint (as defined above) of the first animal within each treatment group, **p = 0.0017 (C), ***p = 0.037 (D) (Log-rank test). (**E, F)** Mouse body weight at the experimental endpoint relative to baseline weights for each treatment group for mice carrying either COG-N-415x (E) or COG-N-496x (F) cells. The means of n = 7 mice are shown ± SEM. Significance was determined using a one-way ANOVA with Tukey’s post-test at each experimental endpoint in C and significance was not reached in any case.

A genetically engineered mouse model (GEMM; Alk-F1178SKI/0;Th-MYCNTg/0 mice) of ALK-expressing NB was also employed in which tumours resembling NB develop in the adrenal glands^45^. A significant reduction in tumour volume was observed in the combination drug treated group, compared to lorlatinib alone (Supplementary Fig. 7 A, C). In keeping with the prior PDX study, no signs of toxicity as indicated by weight loss were observed during treatment (Supplementary Fig. 7B). Overall, these studies confirm the high efficacy and tolerability of combined ALK and FGFR inhibition in both PDXs and an orthotopic model of high-risk NB.

### FGRF2 expression correlates with clinical features suggesting that its inhibition might be more broadly applicable to the treatment of NB patients

Analyses of the existing SEQC (r2-SEQC; GSE49710) dataset of NB patients (n=498) showed that FGFR2 expression levels predict prognosis, whereby higher expression levels correlate with an inferior OS and EFS when considering all patients in this dataset (Fig. 6A-B, top panels, p<0.001 and 0.01 for OS and EFS respectively, Supplementary Fig. 8A). Deeper analysis of other clinical variables in the SEQC dataset showed a correlation between FGFR2 expression and OS when separating male and female patients (p<0.05 in both cases), although this was only significant in the case of EFS for females (p<0.05; Fig. 6 A-B). A strong correlation was also observed for patients older than 18 months or in a high-risk group, although the latter was not significant (Fig. 6 A-B, second panel, Supplementary Fig. 8 B-C respectively). Similar trends were observed for MYCN amplified (n=92) and MYCN non-amplified (n=401) patients, whereby FGFR2 expression correlated with reduced OS and EFS for both groups, although this was not of significant for the MYCN amplified group due to the lower number of patients (Fig. 6 A-B). These data were further validated with the TARGET^46^ dataset (n=247), showing that *FGFR2* transcript levels are higher in HR (n=217) or stage 4 patients (n=216; Fig. 6C), as well as those with an unfavourable tumour histology (n=183, where “unfavourable sample histology” is defined based on the histopathological evaluation of tumour samples according to the International Neuroblastoma Pathology Classification (INPC)). According to these data, NB patients, particularly those older than 18 months and/or those with HR disease, may stand to benefit more from an FGFR2-based therapeutic approach.

**Figure 6.**
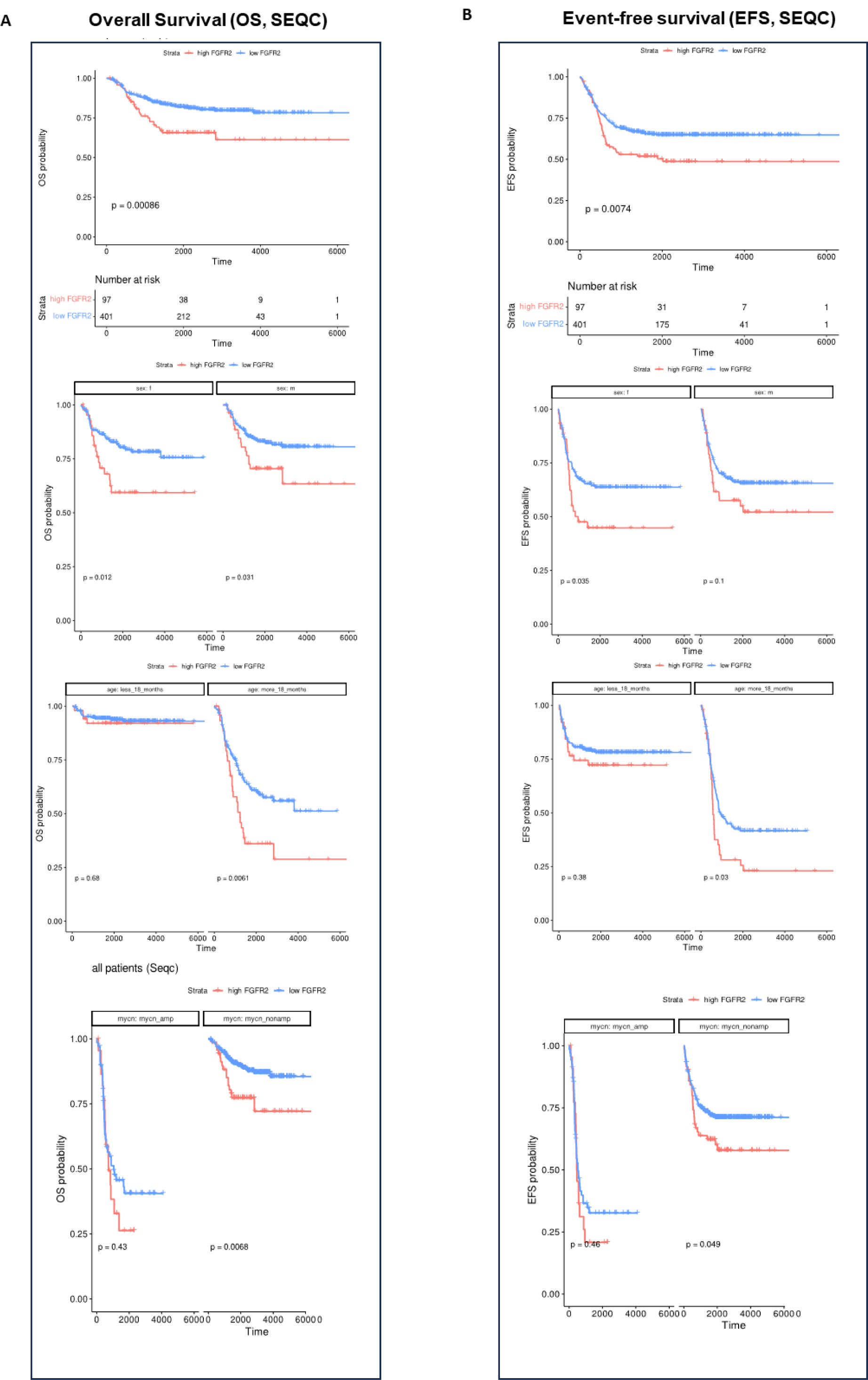
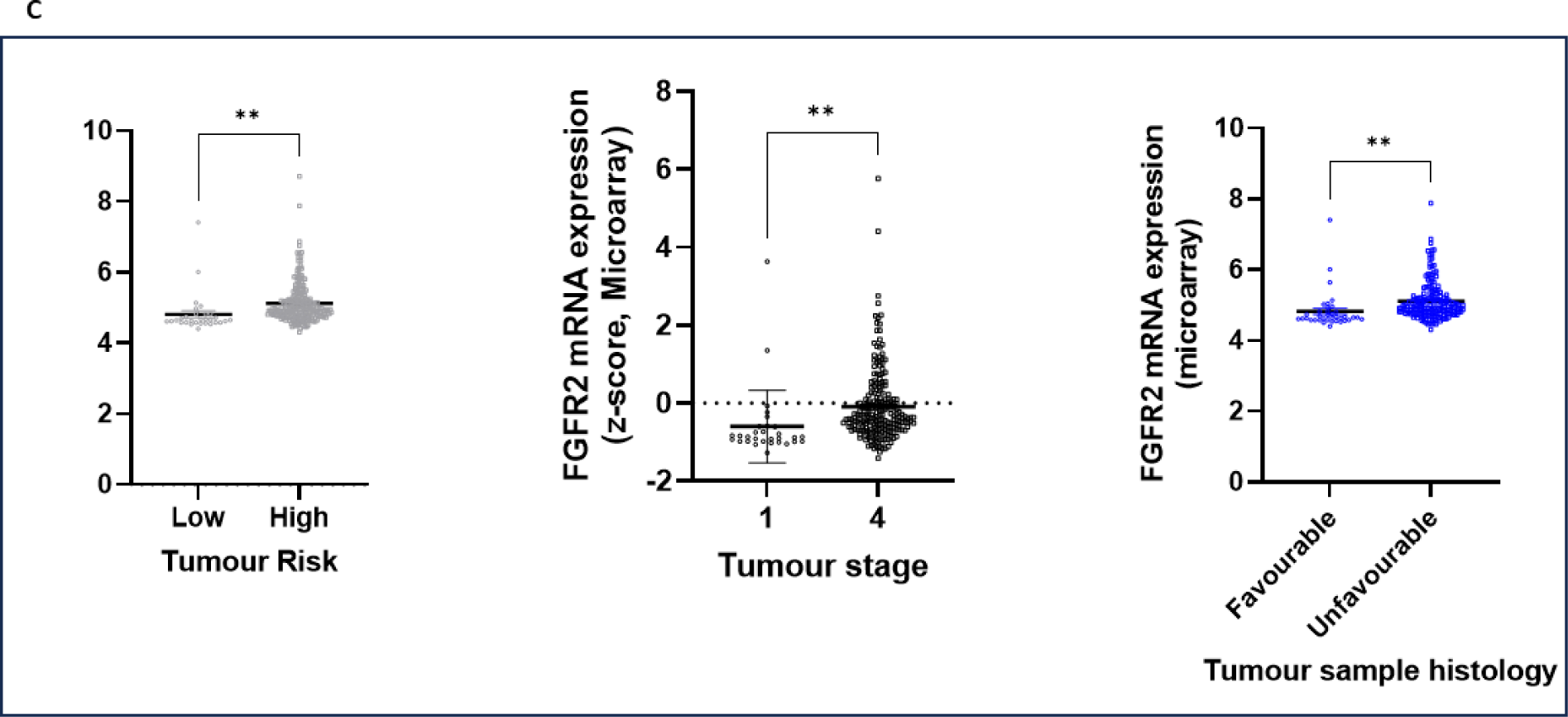
| *FGFR2* mRNA expression correlates with clinical features of NB patients. **(A, B)** *FGFR2* expression is higher in tumours from NB patients with a poorer prognosis, for patients >18 months old and/or those with high-risk disease (expression of FGFR2 mRNA determined from 498 (SEQC) NB patients). Left panels show OS (A) and right panels show EFS (B); all patients are stratified by gender (females or males), age (>18 months or <18 months), or MYCN status (amplification or not amplified). Log-rank test, with p values specified in the figure panels above (significance considered when p<0.05). **(C)** Analyses of the TARGET dataset (n=247), showing FGFR2 transcript levels according to risk group, tumour stage or tumour histology. Significance was determined using an unpaired Student’s t-test with p=0.0060, p=0.0057 and p=0.0031 from left to right, respectively.

## DISCUSSION

NB is the most common extracranial solid cancer diagnosed in children and infants with HR NB cases still lacking effective treatments resulting in a 5-year EFS of less than 50%^6,47^. For these patients, novel therapeutic approaches are urgently needed and while ALK inhibitors, in particular the third-generation ALK TKI lorlatinib, is currently making its way into treatment management for NB patients and showing clinical benefit, resistance/relapse disease is of a distinct probability^3,5^. This issue is not specific to ALK TKIs and has been seen with a variety of similar inhibitors used as monotherapies, representing a recurrent obstacle that hinders their clinical utility, necessitating prolonged treatment courses in many cases. Increasingly, bypass signalling mechanisms of resistance rather than mutation of the target protein-encoding gene are being described, mediated through epigenetic mechanisms circumventing dependence on the target protein for tumour survival^21,24,27,33^. Therefore, it is necessary to investigate resistance bypass mechanisms that arise in response to lorlatinib treatment and to find therapies that target these pathways to extend the efficacy and duration of clinical responses and patient survival.

With this aim, we conducted genome-wide CRISPRa screens in two distinct cell lines to identify genes involved in the lorlatinib response of aberrant ALK-expressing NB; FGFR2 was identified which when upregulated *in vitro* increased NB cell viability in the presence of the ALK TKI lorlatinib. Importantly, a genome-wide CRISPRa screening approach allowed us to fully explore several potential resistance mechanisms in an unbiased manner as we have done previously^,25,48^. Distinct sets of resistance genes were identified across the cell lines, with only three genes (*MET, FGFR2*, and *BACH2*) common to both. Activation of c-MET has previously been shown to confer resistance to alectinib in ALK-positive NSCLC^49^, consistent with a partial functional redundancy of c-MET and ALK due to convergence on several pro-survival signalling pathways^50^. Notably, crizotinib is a potent inhibitor of c-MET^49^ and can overcome ALK inhibitor resistance driven by activation of MET in NSCLC^51^ but has limited efficacy in NB due to the endogenous expression of crizotinib-resistant ALK mutants^52^. BACH2 is a BTB-basic region leucine zipper transcription factor previously shown to have anti-proliferative and anti-differentiation roles in murine NB cells via upregulation of p21 expression^53^. Interestingly, BACH2 serine 510 phosphorylation, which decreases its activity, has been reported to be lower upon treatment with ALK TKIs in NB cells^54^, highlighting a complex link between ALK and BACH2. Another study showed that BACH1, a member of the same family of transcription factors, increases resistance to oxidative stress in SHSY5Y cells^55^, although contradictory evidence exists regarding the anti-proliferative/pro-apoptotic role of these factors^56^. Of more relevance here, both Reactome pathway and GSEA analyses of all significantly enriched genes detected in the CRISPRa screens of both SHSY5Y and CHLA20 cells, showed a prevalence of FGFR2-associated pathways with FGFR2 being represented in all of the top 10 cancer-associated pathways identified. Indeed, overexpression of FGFR2 in SHSY5Y and CHLA20 cells increased the EC_50_ for lorlatinib, and given the availability of inhibitors for this protein, is the focus of this study.

While our results show for the first time the relevance of FGFR2 expression in the context of ALK TKI treatment in NB, upregulation of FGFR2 with subsequent activation of the MEK/ERK pathway and decreased sensitivity to a CHK1 inhibitor has previously been shown in a MYCN amplified, ALK WT NB cell line^57^. Furthermore, another study showed that pharmacological inhibition of FGFR2 overcomes cisplatin resistance more generally in NB via inhibition of apoptosis^58^. These data are in keeping with our observation that FGFR2 is upregulated in LR NB cells compared to their parental counterparts. FGFR proteins activate various signalling pathways, such as MAPK, PI3K, and PLCγ pathways^59^ overlapping with those known to be downstream of ALK activity, suggesting that FGFR2 could relieve dependence on ALK activity in NB, substituting for its activity. Notably, no new mutations in *ALK* were observed in the LR NB cells compared to their parental lines, hence, it is likely that the LR status of the cell lines is due to an ALK-independent mechanism such as up-regulation of FGFR2 and its associated pathways, as our data support.

These data were corroborated with a high-throughput drug screen on parental and LR cell lines using a library of 1500 FDA-approved compounds. In keeping with our prior CRISPRa screen and RNA-Seq data, tyrosine kinase receptors were a common target among drugs that were significantly more efficacious in the LR than in the parental cell lines, including inhibitors targeting the FGFR pathway. High throughput drug screens provide an unbiased drug discovery approach for identifying candidates that exhibit desired anticancer activity without requiring *a priori* knowledge of specific targets that drive cancer progression, and have been previously performed using NB cell lines^60,61^, but not with an FDA-approved drug panel of this size. The screen we conducted identified drugs that have not previously been described as therapeutics for NB, whether ALK TKI sensitive or not. However, several of the drugs identified by this screen have both on-target or off-target effects against multiple tyrosine kinase receptors making it difficult to conclude the exact target in the NB cell lines for each drug, particularly as they are all screened at a standard concentration of 1000nM. Regardless, the predominance of drugs targeting kinase receptors detected in this screen indicate the potential of using a range of TKIs to overcome or prevent lorlatinib resistance in NB.

Given these data, we selected ponatinib for further validation as it targets several kinases as well as FGFR, including PDGFR, SRC, RET, c-KIT, and FLT1^62^. Ponatinib has been approved by the FDA primarily for the treatment of chronic myeloid leukaemia and Philadelphia chromosome-positive acute lymphoblastic leukaemia (Ph+ ALL) as a later-in-line treatment strategy, used in patients that are resistant to at least 2 prior TKIs^62^. However, as ponatinib is not specific to FGFR we also tested the efficacy of erdafitinib, a more selective FGFR inhibitor which is an FDA-approved drug for the treatment of urothelial cancer with FGFR mutations, and is also being investigated as a treatment for other FGFR-driven cancers, including NSCLC and prostate cancer^63^. Indeed, the RAGNAR phase 2 trial is assessing erdafitinib efficacy for the treatment of multiple solid tumours including adult and childhood malignancies, results of which could potentially accelerate its clinical use for NB treatment as proposed here^64,65^. Both ponatinib or erdafitinib in combination with lorlatinib showed synergism at a range of concentrations in both parental and LR cell lines, indicating that these combinations have the potential to prevent or overcome lorlatinib resistance combatting bypass signalling^21,33,66^. Furthermore, the high level of synergy between these drugs indicates that the dose of each drug may potentially be lowered substantially, which will be beneficial in reducing the side-effects associated with treatment, improving the overall quality of life of patients.

Ponatinib has previously been tested for clinical efficacy in combination with other TKIs including trametinib, a MEK inhibitor used in the treatment of KRAS-mutant NSCLC, which served to prevent the emergence of trametinib resistance brought about by FGFR signalling, although this clinical trial reported that the combination of ponatinib and trametinib resulted in substantial toxicities^67^. However, ponatinib has previously been used in combination with asciminib, a BCR-ABL inhibitor, for a Ph+ ALL relapse patient, and remission with manageable side-effects was achieved, highlighting the unpredictable nature of combinatorial regimens using multi-kinase inhibitors^68^. Indeed, in our hands, the combination of FGFR and ALK TKIs was not toxic to the PDX mice when administered at the doses used in our study, while also inducing a significant delay in tumour growth with a prolonged EFS compared to single agent treatment. Notably, one of the PDXs we tested harbours a common ALK mutation (F1174L) as well as having MYCN amplification, representative of aggressive HR NB, and the second is WT for ALK and has MYCN amplification, suggesting this approach may benefit patients with NB of different genetic backgrounds. Further studies of an orthotopic GEMM model of human F1174S ALK and MYCN amplification-driven NB confirmed these findings. Together, these data along with the outcome-predictive significance of FGFR expression levels in patients gleaned from the SEQC and TARGET datasets, show the potential efficacy of combining erdafitinib with lorlatinib for a range of genetically diverse NB, thereby extending the clinical applicability of this therapeutic approach.

Overall, research into cancer drug development has begun to focus on the use of combinatorial targeted agent-based therapies in recognition of the almost inevitable consequences of using single agents, largely resistance^69,70^. Synergistic combinations are predicted to desecrate the tumour in shorter time-frames due to amplified efficacies, allowing lower doses of each drug to be used due to the increase in potencies^71^. Such approaches may also apply in aberrant-ALK expressing NB whereby, as shown here and previously by us and others^24–26^, combined use of inhibitors of defined bypass signalling pathways such as FGFR and ALK may result in better outcomes for patients.

## METHODS

In all results as described in the main text and in the figure legends, biological replicates are intended as cell flasks cultured separately for all *in vitro* experiments. Biological replicates are intended as separate animals in the *in vivo* studies.

### Cell lines and cell culture

The neuroblastoma cell lines, CHLA-20 and COG-N-415 were obtained from the Children’s Oncology Group Childhood Cancer Repository. KELLY and SH-SY5Y cell lines were obtained from the European Collection of Authenticated Cell Cultures. 293FT cells were obtained from Thermo Fisher Scientific. The KELLY-LR and CHLA-20-LR cell lines were generated by initially culturing the parental cells under an inhibitory concentration (IC) 90% (IC90) (10,000nM) for 24 hours followed by the EC_50_ (1000nM). Every 5 passages the concentration of lorlatinib was doubled until 4000nM was reached. CLB-GA and CLB-GA LR cell lines were sourced from Prof Carlo Gambacorti-Passerini and Dr Luca Mologni, University of Milano-Biccoca, Milan, Italy^21^.

CHLA-20, CHLA-20-LR and COG-N-415 cells were cultured in IMDM (Gibco, #21980032) supplemented with 20% FBS, 1% insulin-transferrin-selenium (ITS; Gibco, Cat#41400045) and 1% penicillin and streptomycin (PS). KELLY, KELLY-LR, CLBGA and CLBGA-LR cells were cultured in RPMI 1640 medium (Gibco, Cat#21875091) supplemented with 10% FBS, and 1% PS. SH-SY5Y and 293FT cells were cultured in DMEM (Gibco, Cat#41966029) supplemented with 10% FBS and 1% PS. Cells were grown at 37°C in a humidified incubator with 5% CO_2_. All cells were mycoplasma-free and subjected to quarterly in-house testing.

### Cloning of guide sequences

Double-stranded sequences were generated by phosphorylating and annealing oligonucleotides with T4 polynucleotide kinase (NEB, Cat#M0201) and T4 DNA ligase (NEB, Cat#M0202), respectively. Guide sequences were cloned into the lenti sgRNA(MS2)_puro backbone (Addgene, Cat# 73795) by Golden Gate assembly using BsmBI (NEB, Cat#R0580) and amplified in NEB Stable competent E. coli cells. A full protocol is available at www.sam.genome-engineering.org/protocols.

### Production of lentivirus and cell transduction

Low passage 293FT cells were cultured in DMEM with 5% FBS. Cells were seeded at 40% confluency in six-well plates for small-scale production, or 15 cm dishes for larger-scale production. Once confluency reached 80–90%, cells were transfected with pMD2.G (Addgene, Cat#12259), psPAX2 (Addgene, Cat#12260) and transfer plasmid at equimolar ratios using TransIT-293 transfection reagent (Mirus Bio, Cat#MIR2704) according to the manufacturer’s instructions. The media containing lentivirus was collected after 60 h. Cellular debris was removed following centrifugation at 500 × g and the media was passed through a 0.45 µm polyethersulfone (PES) filter. The supernatant was stored at −80 °C. To generate stable cell lines, cells were seeded into culture vessels in media supplemented with 5% FBS, grown to near-confluency and then transduced with lentivirus-containing media. After 20 h, cells were split into media containing antibiotics as follows: 10 µg/mL blasticidin for 7 days, 300 µg/mL hygromycin for 7 days and 1.5 µg/mL puromycin for 5 days.

### Generation of CRISPR SAM cell lines

CHLA-20 and SH-SY5Y cells were co-transduced with lentiviruses carrying the dCas9-VP64_blast (Addgene, Cat#61425) and lenti MS2-p65-HSF1_hygro (Addgene, Cat#61426) constructs at a multiplicity of infection (MOI)∼0.2 and selected as described above. To confirm CRISPR SAM activity, cells were transduced with lentivirus carrying the lenti sgRNA(MS2)_puro construct (Addgene, Cat# 73795) into which guide sequences were cloned and overexpression of the target genes was confirmed by RT-qPCR.

### CRISPR SAM screens

The puromycin-resistant pooled gRNA library v1 (Addgene, Cat#1000000057) was packaged into lentivirus in 15 cm plates, as described above. CHLA-20 and SH-SY5Y cells stably expressing dCas9-VP64 and MS2-p65-HSF1 were seeded into 15 cm plates and transduced with the lentiviral gRNA library at MOI 0.3–0.4, with a minimum representation of 500 transduced cells per guide. After 20 h, cells were split into 500 µg/mL zeocin for 5 days and passaged every second day, maintaining >500 cells per guide. After 5 days of selection, 3.6 × 10^7^ cells were harvested as a day 0 sample. The remaining cells were split into duplicate populations and cultured in the presence of vehicle (0.3% DMSO) or lorlatinib at EC_50_ –equivalent concentrations as determined by 72-h dose– response assays. Cells were maintained below 80% confluency and harvested after 14 days of treatment for genomic DNA extraction.

### Preparation of HiSeq libraries

Genomic DNA was extracted with the QIAamp DNA Blood Maxi Kit (Qiagen, Cat#51194). The gRNA regions were PCR-amplified for 22 cycles with Herculase II Fusion DNA Polymerase (Agilent, Cat#600677) in 23 replicate reactions, with a total input of 230 µg DNA corresponding to 500X library coverage. Each 100 µL reaction comprised Herculase II reaction buffer (2 mM Mg2+), 1 mM dNTPs, 2% DMSO, 250 nM pooled forward primer, 250 nM specific reverse primer and 10 µg template DNA. The Illumina-compatible primers contained P5/P7 adapters, a staggered region (forward primers only) and 8 bp index barcodes (Supplementary Table 8). PCR products were pooled and precipitated with 400 µL ethanol, resuspended in 100 µL water and resolved on a 3% agarose gel at 100 V for 5 h. The 270–280 bp amplicon was isolated with the QIAquick Gel Extraction Kit (Qiagen, Cat#28706) according to the manufacturer’s protocol, except the QC buffer was incubated at room temperature. Products were tested for concentration and specificity using High sensitivity D1000 ScreenTape and qPCR using the KAPA Library Quantification Kit, multiplexed, spiked with PhiX Control v3 library (Illumina, Cat#FC-110-3001) and run on an Illumina HiSeq 2500 on Rapid Run 1 × 100 bp mode for 115 cycles.

### Gene enrichment analyses

For NGS gene enrichment analysis, raw FASTQ files were downloaded from the Bauer Core server using FileZilla. Initial QC was conducted with FastQC (Babraham Bioinformatics, Cambridge, UK) to assess the general quality of the sequence runs. Raw FASTQ sequencing files were demultiplexed with bcl2fastq2 v2.2, then matched to the guide sequences from the library files using the MAGeCK count function. Read counts per gRNA were calculated by averaging gRNA read counts of two biological replicates per screen condition and normalizing to the total gRNA read count. Normalized read counts in ALK inhibitor-treated cell populations were log-transformed and compared with those in DMSO-treated cell populations to identify gRNAs that were preferentially enriched under ALK inhibitor conditions. An arbitrary threshold beta score >1 was applied and genes with multiple gRNAs exceeding this threshold were considered to be enriched. To improve stringency, only genes enriched in lorlatinib screens with p<0.01 were considered as candidates for downstream analysis.

For Gene set enrichment analysis (GSEA), hallmark and gene ontology gene expression datasets were downloaded from the molecular Signatures database (MSigDB) v6.227 and analyzed with GSEA v3.0 (www.broadinstitute.org/gsea). Enrichment was carried out by calculating overlaps between MSigDB datasets and putative resistance genes.

### Cell viability assays

Cells (10^4^) were seeded into black, clear-bottom Nunc 96, 384 black well optical plates (ThermoScientific) or 96-well White Polystyrene Microplates (Corning, Cat# CLS3610) and after 24 h, the media was aspirated before treatment was carried out with compounds and concentrations as specified in each figure and figure legend. The DMSO concentration was maintained below 0.06% for all conditions. For analysis of cell viability, 20 µL (for 96-well plates) or 5µL (for 384-well plates) of CellTiter-Blue reagent (Promega, Cat# G8080) was added to the cells, according to the manufacturer’s protocol. Fluorescence was read on a SpectraMax i3 microplate reader (Molecular Devices).

### RNA-Sequencing and analysis

RNA-Seq was performed at Novogene (UK) with RNA-Seq library preparation and quality control for subsequent RNA-Seq using the Illumina NovaSeq 6000. Compressed FASTQ files were retrieved from the sequencing facility and assessed for quality with FASTQC (v0.11.9). Transcript abundances were estimated with Salmon (v1.9.0). Differential expression testing was performed in R (v4.1) using the tximeta (v1.12.0) and DESeq2 (v1.34.0) packages, with cell line identity included as a covariate. Significantly differentially expressed genes were then defined as those for which the adjusted *p*-value was less than 0.05 and the absolute log₂ fold change was greater than 1.5. To call single nucleotide variants, reads were first trimmed to remove sequencing adapter contamination using trim-galore (v0.6.7) with default settings. The trimmed reads were then aligned to the human genome (GRCh38.p14) in a splice-aware manner using STAR (v2.7.1a). Variant calling was performed according to the GATK best practices workflow for small RNA variants using GATK tools (v4.2.6.1). Fusion gene detection was performed with STAR-FUSION (v1.10.0).

### High-Throughput Drug screens

The FDA-Approved Drug Screen Library (Selleckchem) of 1430 drugs, was utilised for high-throughput drug screens. The FDA library was supplied as a collection of 20 ‘Master’ plates in a 96-well format. Using a Beckman Coulter Biomek® NXP laboratory automation workstation, these drugs were added to the cells at a final concentration of 1000nM.

KELLY, KELLY-LR, CHLA-20 or CHLA20-LR cells (3000-9000 cells/well in 45μL of appropriate media) were seeded in clear bottom Nunc 384-well optical plates (ThermoFisher). After a 24-hour incubation at 37°C, 5% CO_2_, 5μL of drugs from the drug library were added to bring the final volume to 50μL. Plates were then incubated for a further 72-hours under the same conditions. All drugs were applied to the cells at a final concentration of 1000nM before plates were covered with breathable AeraSealTM sheets (Alpha Laboratories) to prevent evaporation. At the end of the 72-hours, fluorescence was measured on a SpectraMax i3 microplate reader (Molecular Devices).

The fluorescence of each well was normalised to the median fluorescence of cells treated with 0.01% DMSO. Drug screen data analysis was performed using packages within R studio (v4.1). Analysis was divided into two parts: quality control and identification of efficacious compounds. R studio packages were used throughout the analysis for data visualisation, manipulation and labelling, including tidyverse (v1.3.1) and ggrepel (v0.9.1). To assess the effect of compounds on cell viability, the median percentage viability of technical replicates associated with the treatment of each compound was used. The log ratio of viability between the parental and corresponding LR cell line was calculated and drugs with a log ratio greater than 0.7 in both cell lines were identified. The drug targets and pathways associated with the isolated drugs were also identified and visualised using bubble plots created by the R studio package packcircles (v0.3.4). The drug target and pathway data of the FDA-approved drug library were provided by SelleckChem.

### Synergy experiments

For dose-response matrices, cells were treated with log-scale concentrations of lorlatinib in addition to log-scale concentrations of ponatinib or erdafitinib, as indicated in 7×7 grids, and the DMSO concentration was maintained below 0.06%. Potential synergy between the compounds was evaluated by calculating the synergy score based on the Loewe model^72^ using synergy finder^73^. The synergy score was calculated for each combination of drug concentrations and also as an overall value and is defined as: >10 = synergistic; Between –10 and 10 = additive; <-10 = antagonistic.

### RNA extraction and RT-qPCR analysis

Total RNA was isolated from cell lines using the RNeasy Plus Mini Kit (Qiagen, Cat#74134). In total, 1 μg RNA was reverse transcribed using the High-Capacity cDNA Reverse Transcription Kit (Thermo Fisher Scientific cat# 4368814) followed by the TaqMan RT-qPCR assay. Relative quantification (ΔΔC_T_) analysis was conducted with normalization to GAPDH (Thermo Fisher Scientific, Hs02786624_g1, 4331182) and HPRT1 (Thermo Fisher Scientific, Hs02800695_m1, 4331182). All reactions were performed in technical triplicates.

### Immunoblot analysis

Adherent cells were washed, and lysates prepared with prechilled RIPA Lysis and Extraction Buffer (Thermo Fisher Scientific, Cat#89900) supplemented with 1% Halt Protease and Phosphatase Inhibitor Cocktail (Thermo Fisher Scientific, Cat#78440). Around 30 μg of protein lysate per sample was resolved by SDS-PAGE and transferred to a PVDF membrane with the Semi-Dry and Rapid Blotting System according to the manufacturer’s protocol (Bio-Rad). The membrane was blocked in 5% BSA and incubated with antibodies overnight at 4°C. Primary antibodies were as follows and diluted to 1:1000 unless specified otherwise: anti-FGFR2 (CST, cat# 23328), anti-β-Tubulin (CST, cat#2146). Membranes were then incubated with secondary antibodies for 1 h at room temperature (1:10,000 anti-rabbit HRP-immunoglobulins (Dako, Cat#P0448)). If required, membranes were stripped once with stripping buffer (Thermo Scientific™ Cat# 46430) and re-blocked. Membranes were developed with Immobilon Western Chemiluminescent HRP Substrate (Merck Millipore), and bands imaged with an ImageQuant LAS-4000 imaging system (GE Life Sciences).

### Patient-derived xenograft studies

NSG mice were obtained from Charles River and housed in groups of 2–6 mice per cage. All procedures were carried out under UK Home Office licence P4DBEFF63 according to the Animals (Scientific Procedures) Act 1986 and were approved by the University of Cambridge Animal Welfare and Ethical Review Board (AWERB). COG-N-496x and COG-N-415x patient-derived xenograft (PDX) cells were obtained from the Childhood Cancer Repository maintained by the Children’s Oncology Group (COG). Cells were suspended in Matrigel (Corning) diluted 1:2 with PBS and 5 × 10^5^ cells (300 µL) were injected into the left flank of NSG mice at on average 8 weeks of age. Tumours were measured daily with manual callipers and tumour volumes estimated using the modified ellipsoid formula: V = ab^2^/2, where a and b (a > b) are length and width measurements respectively. Once tumours reached approximately 100 mm^3^, mice were randomly allocated into four treatment groups (n = 6-8 per group) and treated daily with the following agents by oral gavage at 10 µL/g body weight: vehicle (20% hydroxypropyl-beta cyclodextrin), erdafitinib (30 mg/kg), lorlatinib (10 mg/kg) or a combination of erdafitinib and lorlatinib at the same doses. Mice were euthanized once tumours reached 15 mm in any direction (defined as an endpoint event for EFS analysis).

### *In vivo* studies with GEMM models and ultrasound imaging

All GEMM animal work was carried out by trained personnel and according to Gothenburg Animal Ethics Committee approval, Jordbruksverket (1890–2018, 3225–2020, 4793-2023). Alk-F1178S KI/0;Th-MYCN Tg/0 (–backcrossed onto a 129×1/SvJ background^45^) mice were dosed by oral gavage once daily with erdafitinib 30mg/kg (MedChemExpress, HY-18708/CS-4988) and/or lorlatinib 10mg/kg (Reagency, RLor-10-21) dissolved in DMSO (Sigma-Aldrich, #D4540), PEG 300 (Sigma-Aldrich, 90878) and H_2_0. A Vevo 3100 imaging system (FUJIFILM VisualSonics) was used to screen GEMM 2-3 times a week. A 3D scan was performed when the tumour reached a volume of 210-260mm^3^ after which treatment was initiated (Day 0) with either erdafitinib 30mg/kg + lorlatinib 10mg/kg (n=3) or lorlatinib 10mg/kg (n=3). Volume calculations were performed for Day 0, Day 7 and Day 14 tumours from data either obtained by Vevo 3100 imaging systems employing VevoLab (FUJIFILM VisualSonics) 3D Multi-Slice Method Open, measuring the diameter in three dimensions in VevoLab, or by direct measurements of the diameter in three dimensions at sacrifice. For the tumours where the diameter in three dimensions was obtained, the volume was calculated by employing the equation for the volume of an ellipsoid 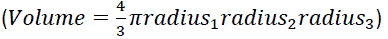. 2D images were acquired in VevoLab, exported in a TIF format and cropped into Affinity Designer (Serif, 1.10.0.1127).

### Patient data analysis

CBioPortal (www.cbioportal.org) was queried using a publicly available TARGET study (TARGET, 2018, phs000218 (https://ocg.cancer.gov/programs/target) available at https://portal.gdc.cancer.gov/projects, Study ID phs000467. This dataset consists of 1,089 samples from 1,076 patients. Among them, 143 samples with bulk RNA-Seq RPKM data, 249 with Agilent microarray and 59 with copy number alterations are available. The SEQC and Kocak datasets from the platform R2 were also analysed. By default, the R2 platform “scans” a range of cutoff values, applying a log-rank test to each, and selects the cut-off point which yields the minimum p-value. Using this methodology, type I error is inflated due to hundreds of comparisons, necessitating multiple-testing correction. In our analyses, we instead manually chose an FGFR2 expression value cut-off, based on the expression distribution of the cohort. This was done because the population distribution appears bimodal.

### Statistical analysis

All Student’s *t*-tests, One– and Two-way ANOVA models, correlation analyses and Kaplan-Meier survival analyses were conducted with GraphPad Prism 9/10 software.

## Acknowledgements

This research was supported by the Cambridge NIHR BRC Cell Phenotyping Hub. J.K. acknowledges support from the NIHR Cambridge Biomedical Research Centre. We would like to acknowledge the Childhood Cancer Repository who provided the neuroblastoma PDX (Texas Tech University Health Sciences Center School of Medicine, Lubbock, TX 79430, USA) and are funded by Alex’s Lemonade Stand Foundation for Childhood Cancer and Prof Carlo Gambacorti-Passerini and Dr Luca Mologni, University of Milano-Biccoca (Milan, Italy) for provision of the CLBGA and CLBGA-LR cell lines.

## Author Contributions

Conceptualization: PP, SDT

Methodology: PP

Investigation: PP, CB, RT, LJ, MB, JDM, CS, LH, EJ, MS, NP, LK, GAAB, BH, RP

Funding acquisition: SDT

Resources: SDT

Supervision: SDT

Writing: PP, SDT

All authors approved the final version of the manuscript.

