## Supplemental Figures for "FGFR2 promotes resistance to ALK tyrosine kinase inhibitors and its inhibition acts synergistically with lorlatinib in the treatment of ALK-expressing neuroblastoma"

**Supplementary Figures**

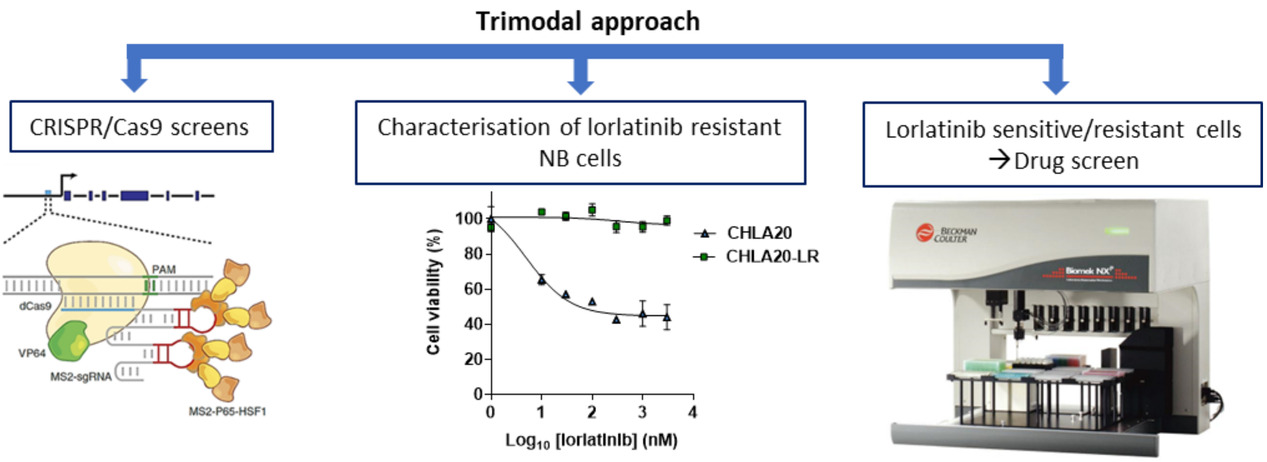


**Supplementary Figure 1 | Overview of the experimental approach used in this study.**

With a trimodal approach consisting of genome-wide CRISPR-dCas9 activation (CRISPRa) screens, RNA sequencing of ALK TKI resistant cell lines and high-throughput drug screens using a library of 1430 FDA approved compounds, we have identified genes whose expression is associated with decreased sensitivity to lorlatinib, creating novel therapeutic vulnerabilities.

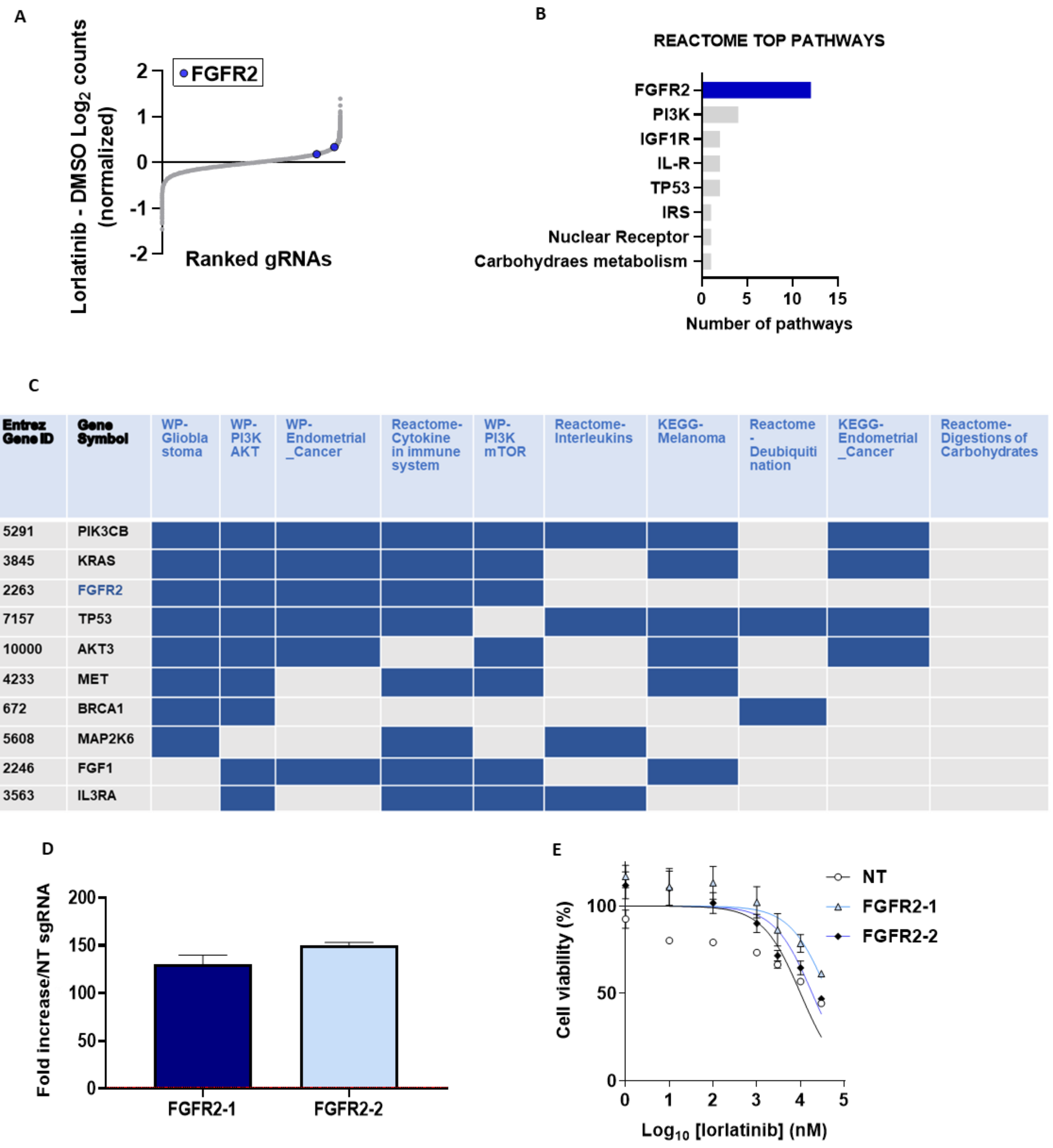


**Supplementary Figure 2 | CRISPRa screens identify FGFR2 expression as a desensitizer to lorlatinib treatment in SHSY5Y NB cells.** (**A**) The gRNAs detected in the SHSY5Y CRISPRa screen are ranked by fold-change vs difference in log-normalized read counts between DMSO and lorlatinib-treated SHSY5Y cells (beta score>1, p<0.01, permutation-based non-parametric analysis (PBNPA)[41]). Two gRNAs per gene are shown and the gRNAs to FGFR2 are highlighted in blue. (**B, C**) Pathway analyses of SHSY5Y CRISPRa screen data: Reactome in B and GSEA with GO gene sets in MSigDB in C. (**D**) FGFR2 expression levels in SHSY5Y-dCas9 cells transduced with one of two independent FGFR2 gRNAs (FGFR2-1 and FGFR2-2) compared to non-targeting (NT) gRNA transduced cells. (**E**) Viability (cell-titer blue (CTB)) of SHSY5Y-dCas9 cells transduced with NT gRNA or two different FGFR2 gRNAs following treatment with the indicated doses of lorlatinib for 72 h. Data points represent the means and SEM of biological duplicates.

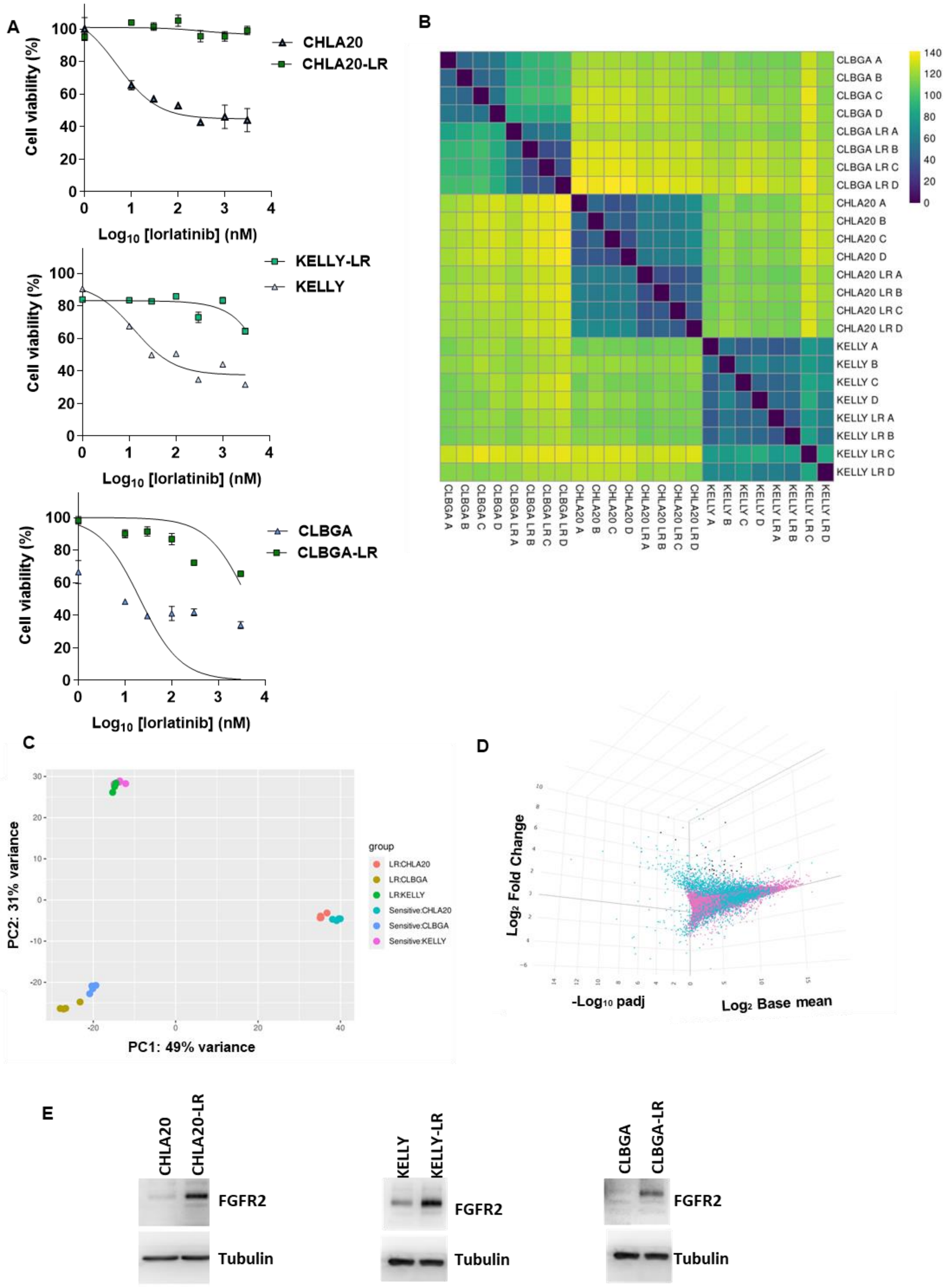
A
Cell viability (%)
Log10 [lorlatinib] (nM)
CHLA20
CHLA20-LR
KELLY-LR
KELLY
CLBGA
CLBGA-LR
B
CLBGA A
CLBGA B
CLBGA C
CLBGA D
CLBGA LR A
CLBGA LR B
CLBGA LR C
CLBGA LR D
CHLA20 A
CHLA20 B
CHLA20 C
CHLA20 D
CHLA20 LR A
CHLA20 LR B
CHLA20 LR C
CHLA20 LR D
KELLY A
KELLY B
KELLY C
KELLY D
KELLY LR A
KELLY LR B
KELLY LR C
KELLY LR D
C
PC2: 31% variance
PC1: 49% variance
group
LR:CHLA20
LR:CLBGA
LR:KELLY
Sensitive:CHLA20
Sensitive:CLBGA
Sensitive:KELLY
D
Log2 Fold Change
-Log10 padj
Log2 Base mean
E
CHLA20
CHLA20-LR
FGFR2
Tubulin
KELLY
KELLY-LR
FGFR2
Tubulin
CLBGA
CLBGA-LR
FGFR2
Tubulin

**Supplementary Figure 3 : Lorlatinib resistant cells (LR) upregulate FGFR2.**

**(A)** Viability of CHLA20, CLBGA, KELLY and their respective LR counterparts (cell-titer blue (CTB)) upon treatment with log scale concentrations of lorlatinib for 72 h. Data represent the means and SEM of two replicates. **(B)** Principal component analysis (PCA) of RNA-Seq data from NB cells (CHLA20, KELLY and CLBGA) and their lorlatinib resistant (LR) counterparts. **(C)** Correlation matrix heatmap showing the Euclidean distance between samples (from dark blue to yellow indicating high to low transcriptome similarity respectively) of RNA-Seq data from NB cells (CHLA20, KELLY and CLBGA) and their LR derivate lines (LR). **(D)** 3D Plot of log2 fold change (FC) against the –log10 of P adjusted values (Adjp) and the log2 base mean of RNA of all lines shown in B. Log2 FC>0 = genes with higher expression in LR cell lines. FGFR2 is in the 14 top upregulated genes (black dots) with a base mean > 500, log2FC> 1.5 and Adjp < 0.05 with base mean >100 within single resistant cell types. **(E)** WB of FGFR2 in CHLA20, CLBGA, KELLY and their respective LR counterparts. Beta-tubulin is used as the reference housekeeping gene. Results shown are representative of two biological replicates.

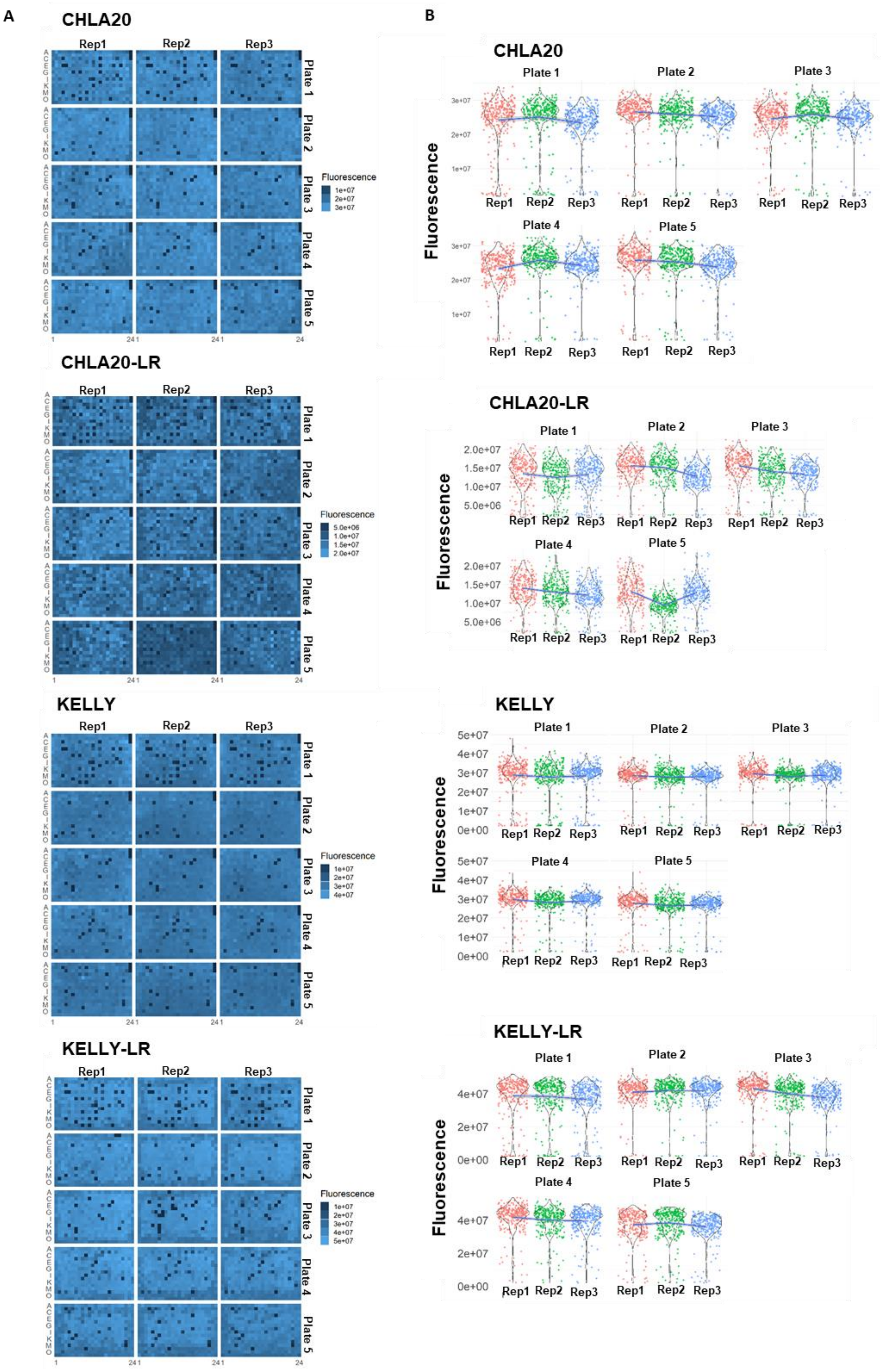
A
B
CHLA20
CHLA20-LR
KELLY
KELLY-LR
Rep1
Rep2
Rep3
Plate 1
Plate 2
Plate 3
Plate 4
Plate 5
Fluorescence
1e+07
2e+07
3e+07
4e+07
5e+07
5.0e+06
1.0e+07
1.5e+07
2.0e+07
0e+00
A
C
E
G
I
K
M
O
1
24

C

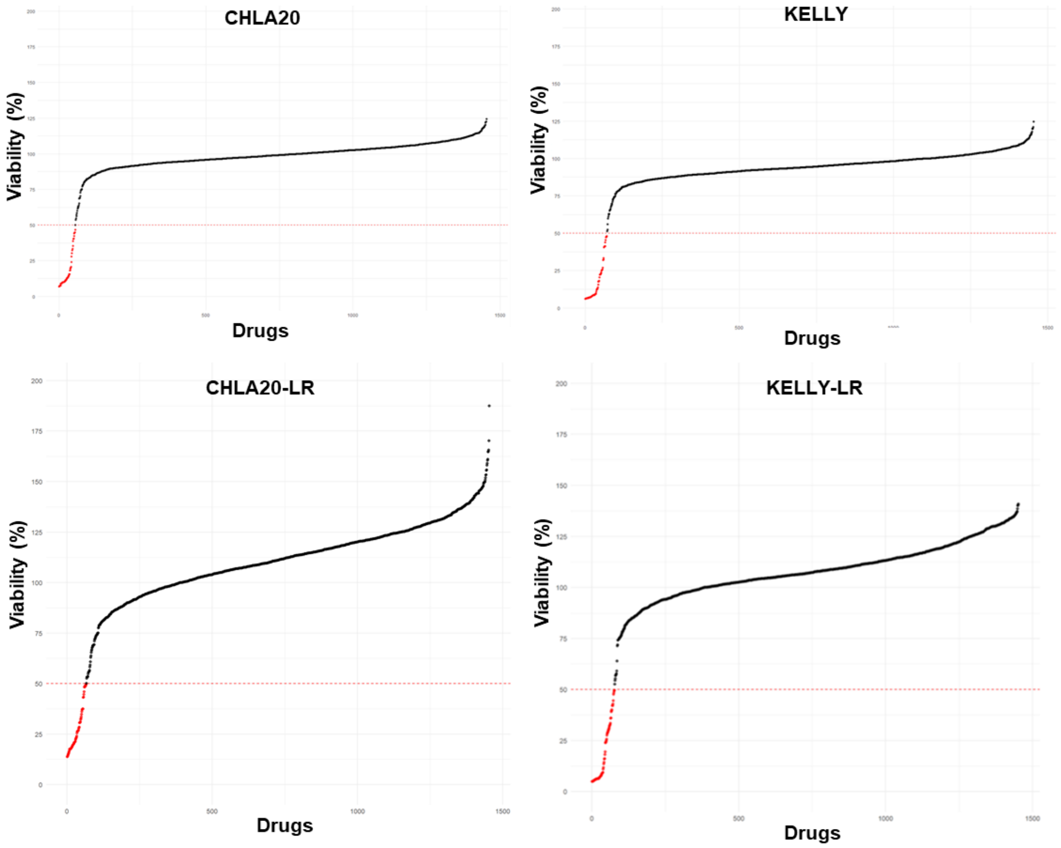

**E**

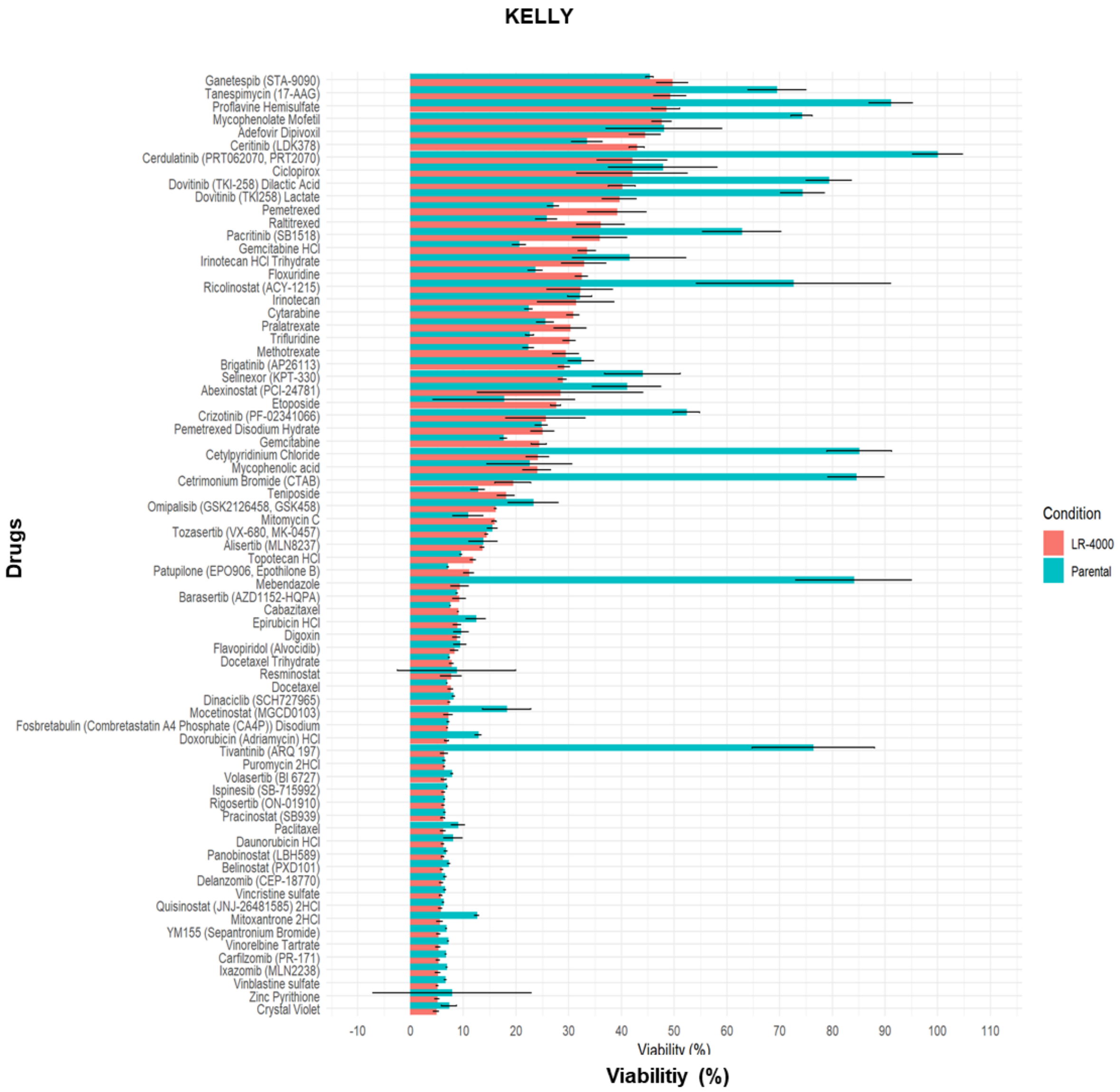
KELLY
Ganetespib (STA-9090)
Tanespimycin (17-AAG)
Proflavine Hemisulfate
Mycophenolate Mofetil
Adefovir Dipivoxil
Ceritinib (LDK378)
Cerdulatinib (PRT062070, PRT2070)
Ciclopirox
Dovitinib (TKI-258) Dilactic Acid
Dovitinib (TKI258) Lactate
Pemetrexed
Raltitrexed
Pacritinib (SB1518)
Gemcitabine HCl
Irinotecan HCl Trihydrate
Floxuridine
Ricolinostat (ACY-1215)
Irinotecan
Cytarabine
Pralatrexate
Trifluridine
Methotrexate
Brigatinib (AP26113)
Selinexor (KPT-330)
Abexinostat (PCI-24781)
Etoposide
Crizotinib (PF-02341066)
Pemetrexed Disodium Hydrate
Gemcitabine
Cetylpyridinium Chloride
Mycophenolic acid
Cetrimonium Bromide (CTAB)
Teniposide
Omipalisib (GSK2126458, GSK458)
Mitomycin C
Tozasertib (VX-680, MK-0457)
Alisertib (MLN8237)
Topotecan HCl
Patupilone (EPO906, Epothilone B)
Mebendazole
Barasertib (AZD1152-HQPA)
Cabazitaxel
Epirubicin HCl
Digoxin
Flavopiridol (Alvocidib)
Docetaxel Trihydrate
Resminostat
Docetaxel
Dinaciclib (SCH727965)
Mocetinostat (MGCD0103)
Fosbretabulin (Combretastatin A4 Phosphate (CA4P)) Disodium
Doxorubicin (Adriamycin) HCl
Tivantinib (ARQ 197)
Puromycin 2HCl
Volasertib (BI 6727)
Ispinesib (SB-715992)
Rigosertib (ON-01910)
Pracinostat (SB939)
Paclitaxel
Daunorubicin HCl
Panobinostat (LBH589)
Belinostat (PXD101)
Delanzomib (CEP-18770)
Vincristine sulfate
Quisinostat (JNJ-26481585) 2HCl
Mitoxantrone 2HCl
YM155 (Sepantronium Bromide)
Vinorelbine Tartrate
Carfilzomib (PR-171)
Ixazomib (MLN2238)
Vinblastine sulfate
Zinc Pyrithione
Crystal Violet
Drugs
-10
0
10
20
30
40
50
60
70
80
90
100
110
Viabilitiy (%)
Condition
LR-4000
Parental

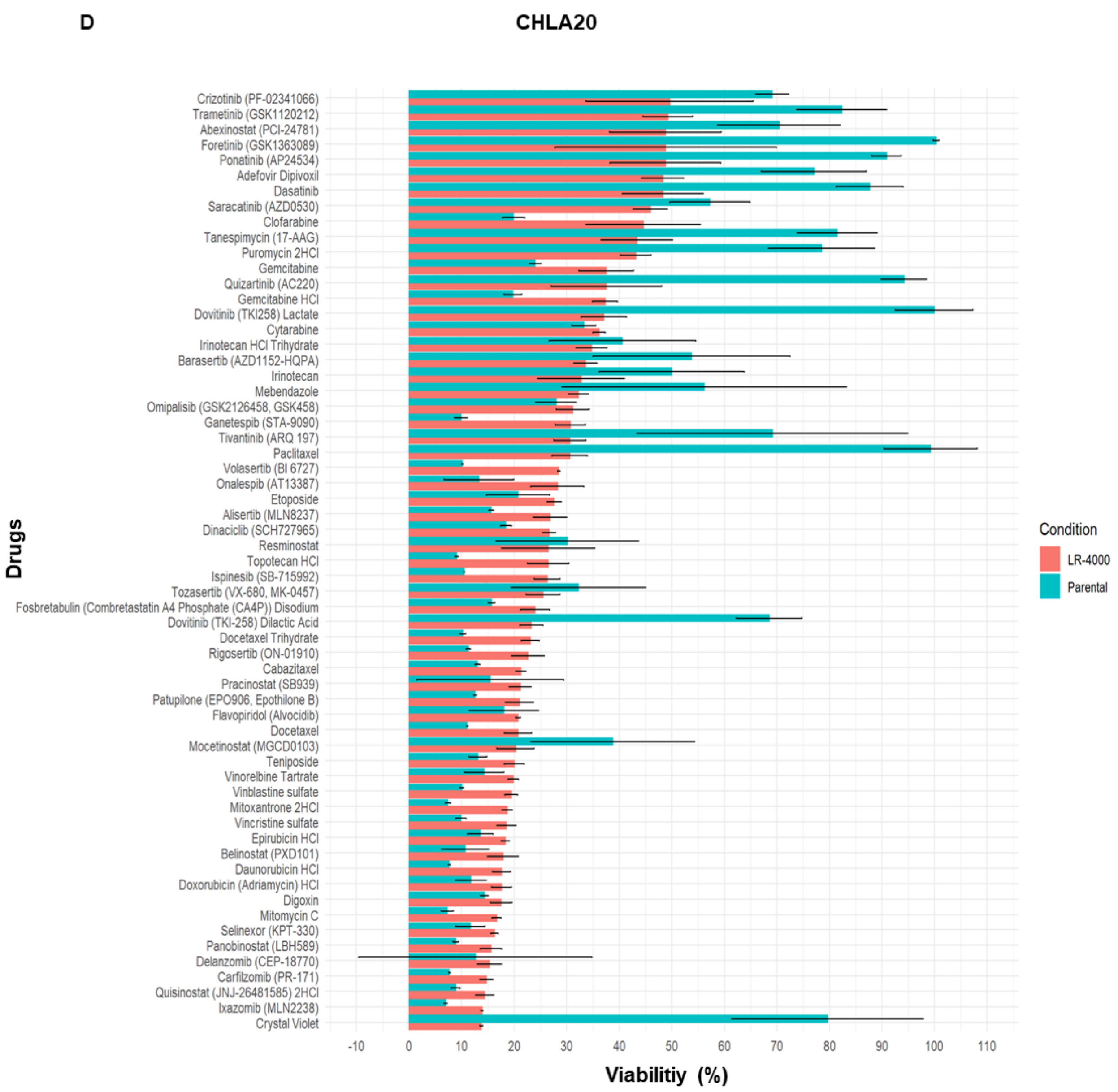


**Supplementary Figure 4 | High-throughput FDA-approved drug screens conducted in CHLA20 and KELLY cell lines, and their LR counterparts.**

**(A)** Fluorescence intensity of individual wells across the fifteen 384-well plates used in the screen showing similar patterns of fluorescence intensity for each plate across the four cell lines. On the left of rep1 of each plate the letters refer to plate rows; along the bottom of plate 5, 1 to 24 refers to columns. **(B)** Violin plots of fluorescence values across plate replicates showing even distribution across plates. The colour within each graph, grouped by drug plate, refers to a different replicate. The width of each violin relates to the frequency of data points at the associated fluorescence value whereby the wider the violin, the higher the density of data points. The horizontal line represents the Locally Weighted Scatterplot Smoothing (LOESS) line. **(C)** The viability of the indicated cell lines following 72-hour exposure to all 1430 compounds used in the drug screens. The red Y intercept and the points illustrated in red highlight the cut off at a reduction of 50% cell viability compared to the DMSO control treated cells. **(D-E)** Drugs that cause a greater than 50% decrease in viability in the LR cell lines (red) but not for the parental cell line (blue). Data represent the median ± standard error of triplicates from different plates.

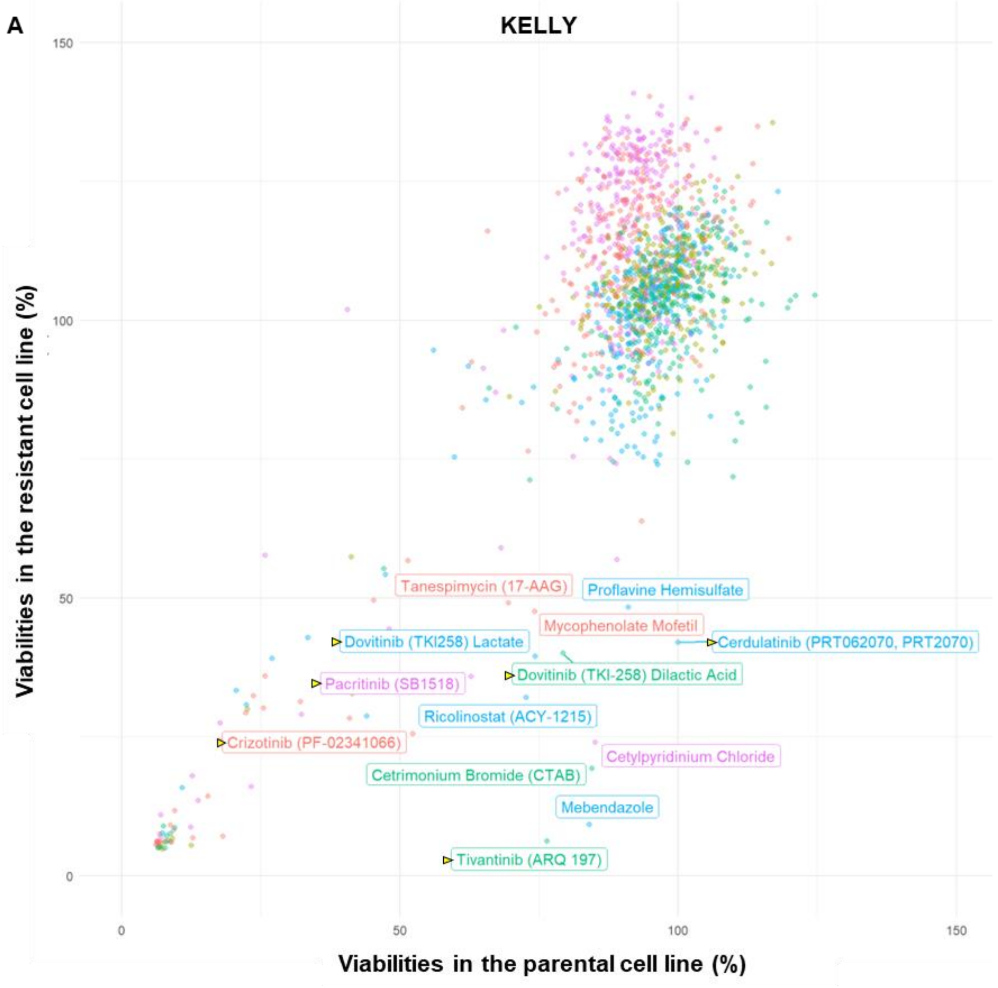

A
KELLY
Viabilities in the resistant cell line (%)
Viabilities in the parental cell line (%)
0
50
100
150
Tanespimycin (17-AAG)
Proflavine Hemisulfate
Mycophenolate Mofetil
Dovitinib (TKI258) Lactate
Cerdulatinib (PRT062070, PRT2070)
Dovitinib (TKI-258) Dilactic Acid
Pacritinib (SB1518)
Ricolinostat (ACY-1215)
Crizotinib (PF-02341066)
Cetylpyridinium Chloride
Cetrimonium Bromide (CTAB)
Mebendazole
Tivantinib (ARQ 197)


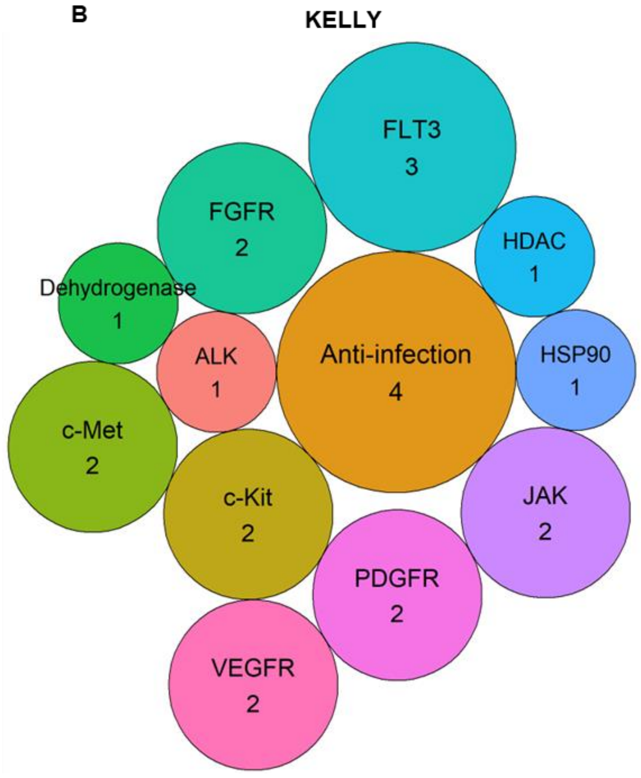

B
KELLY
FLT3
3
FGFR
2
HDAC
1
Dehydrogenase
1
ALK
1
Anti-infection
4
HSP90
1
c-Met
2
c-Kit
2
JAK
2
PDGFR
2
VEGFR
2

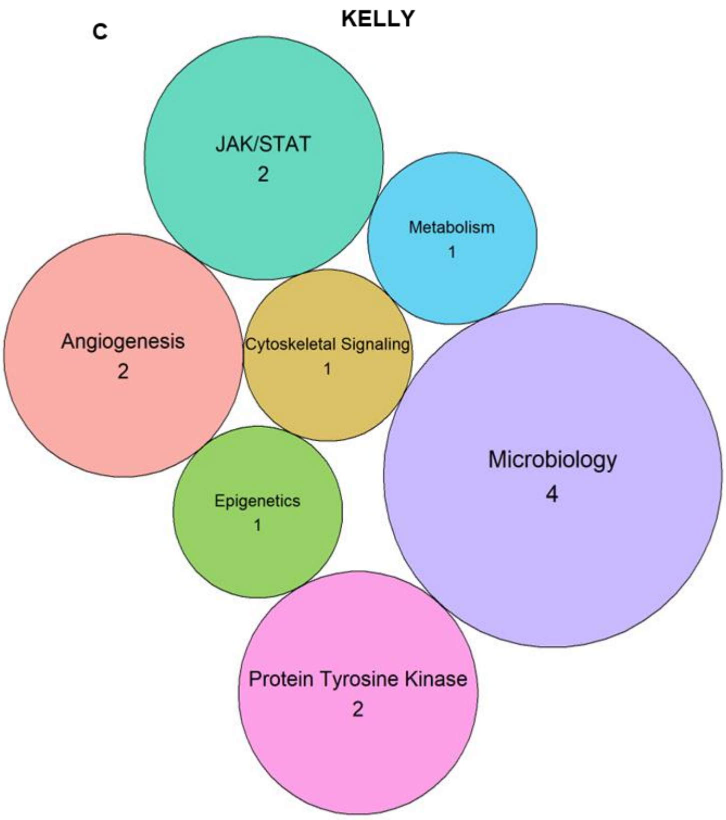


**Supplementary Figure 5 | High-throughput FDA-approved drug screens in KELLY and KELLY-LR cells.**

**(A)** High-throughput FDA-approved drug (~1500 drugs) screens performed with KELLY and KELLY-LR cell lines showing cell viabilities upon drug treatment (all drugs at 1000nM) for 72h. Those highlighted are the drugs that reduced viability of the LR cells with below 50% viability, without similarly affecting the parental cells. The colours represent individual 384-well plates according to where the drugs are located in the library. Yellow triangles next to drug labels highlight the drugs inhibiting kinases. **(B)** Bubble plots showing the common drug targets for drugs more effective in KELLY-LR cells than the parental cell line. **(C)** Bubble plots showing the common families of grouped drugs (including protein tyrosine kinase) for drugs more effective in KELLY-LR cells than the parental cell line.

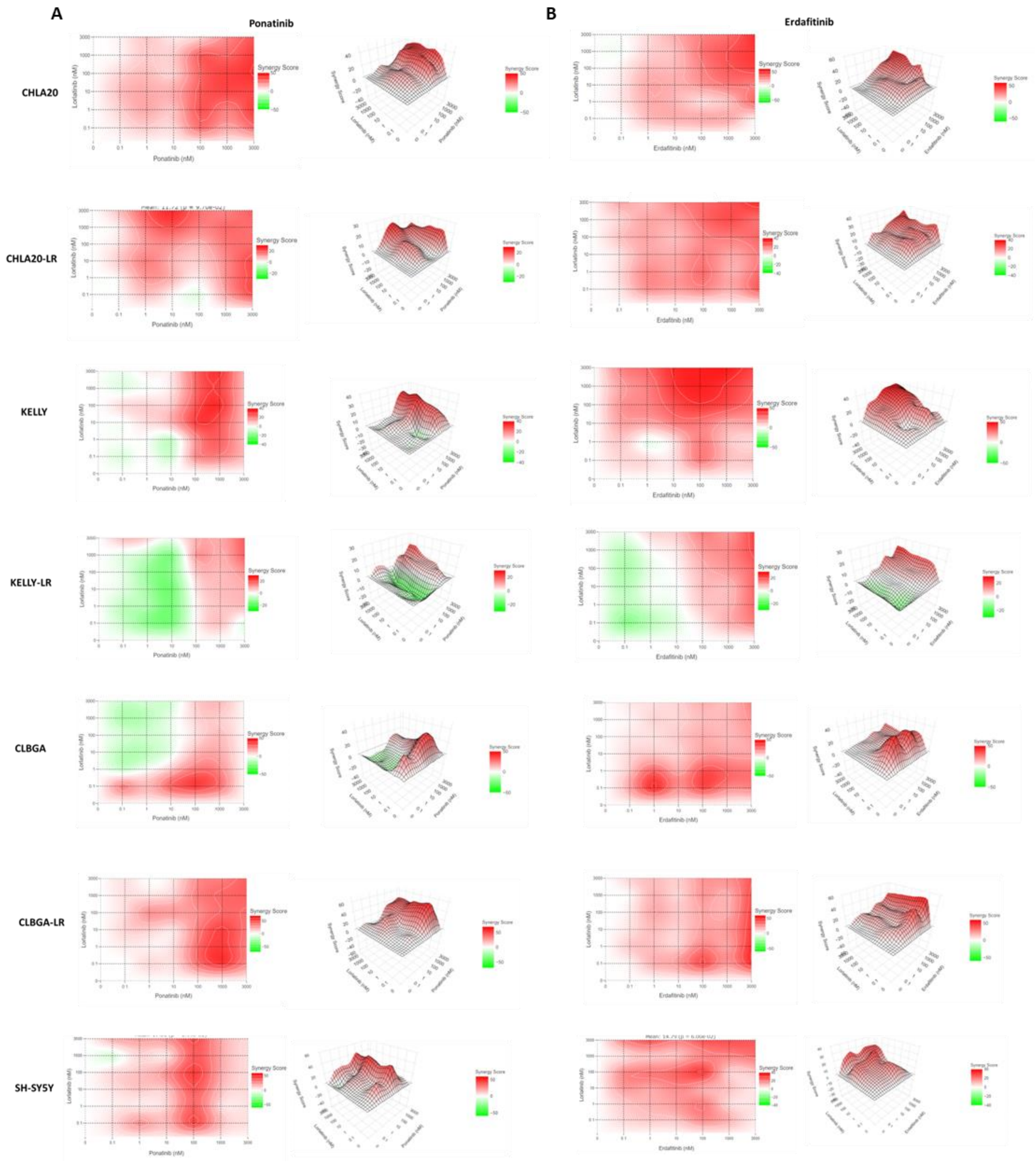


**Supplementary figure 6 | 2D and 3D plots showing synergy scores following pharmacological inhibition of FGFR in combination with lorlatinib in NB cell lines and their LR counterparts.**

**(A)** Two-dimensional and three-dimensional representations of the data shown in Figure 4 (red colour gradient indicates the level of synergy) in NB cells and LR counterparts treated with ponatinib or lorlatinib either alone or in combination at different concentrations (0.1-3000nM for all drugs). **(B)** Two-dimensional and three-dimensional representations of the data shown in Figure 4 (red colour gradient indicates the level of synergy) in NB cells and LR counterparts treated with erdafitinib or lorlatinib either alone or in combination at different concentrations (0.1-3000nM for both drugs). Data shown are from two biological replicates.

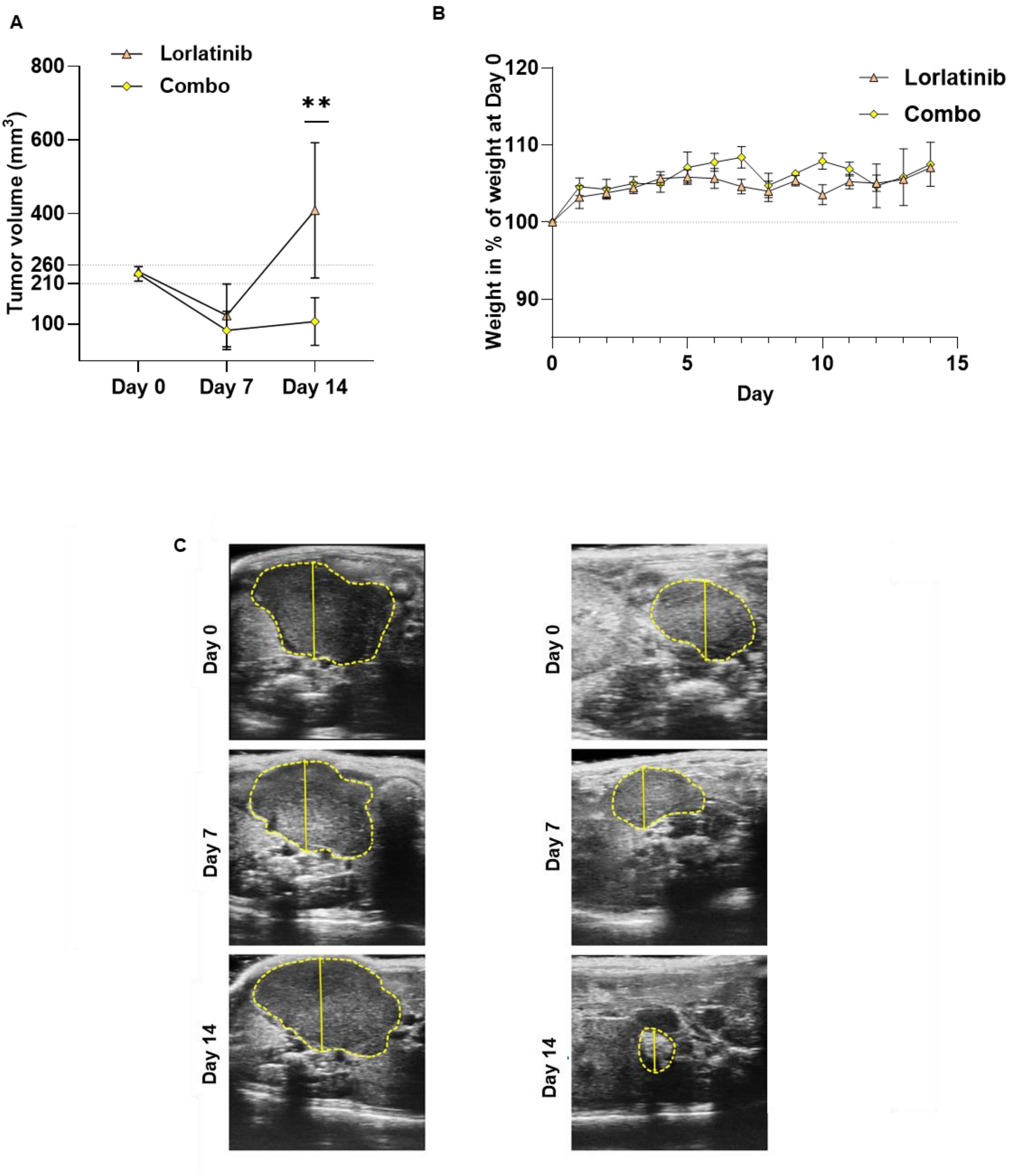


**Supplementary Figure 7 | A combination of lorlatinib with the FGFR inhibitor erdafitinib significantly reduces tumour growth *in vivo* in GEMM models of NB.**

**(A)** Tumour volume over time of Alk-F1178SKI/0;Th-MYCNTg/0 mice treated by daily oral administration of either vehicle (20% hydroxypropyl beta cyclodextrin), lorlatinib (10 mg/kg), or lorlatinib and erdafitinib (30mg/kg) (combo, same doses) for 14 days following treatment initiation. Data shown represent means ± SEM from 3 mice at each time point. Significance was determined using a one-way ANOVA with Tukey's post-test at the experimental endpoint in A. **p = 0.0042. **(B)** Daily mouse body weights recorded until the experimental endpoint relative to baseline weights for each treatment group. Data represent means (n = 3 mice) ± SEM.

Significance was determined using a one-way ANOVA with Tukey's post-test at each experimental endpoint in B and significance was not reached in any case. **(C)** Ultrasound imaging of animals at days 0, 7 and 14 of treatment with lorlatinib (10 mg/kg) or a combination of lorlatinib (10 mg/kg) with erdafitinib (30mg/kg). Magnification bars from top to bottom= 5.73, 5.21, 6.15 mm (left panel); 3.54, 3.49, 2.19 mm (right panel).

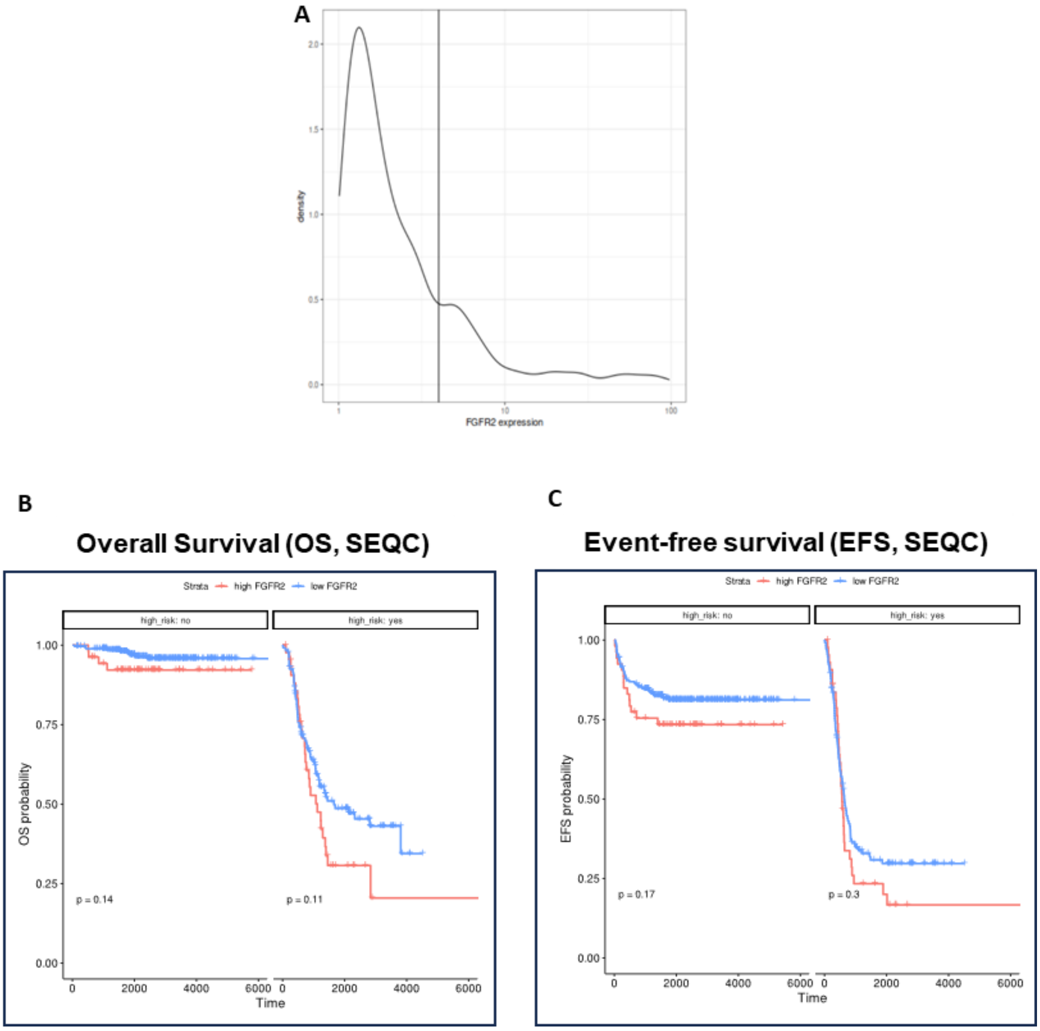


**Supplementary Figure 8 | NB patient survival stratified by FGFR2 expression.** (**A**) Density plot of the patient populations described in the SEQC datasets, according to FGFR2 expression levels. (**B, C**) Expression of FGFR2 mRNA in 498 (SEQC) NB patients according to OS (B) and EFS (C).
